# Integrative nascent RNA methods to reveal cell-type specific transcription programs in peripheral blood and its derivative cells

**DOI:** 10.1101/836841

**Authors:** Samantha Sae-Young Kim, Eunyoo Kim, Aileen Kang Kimm, Hojoong Kwak

## Abstract

Nascent RNA sequencing is a powerful method to measure transcription with high resolution, sensitivity, and directional information, which gives distinctive information about transcription from other methods such as chromatin immunoprecipitation or mRNA sequencing. We present an integrated package of nascent RNA-seq methods - ultrafast Precision Run On (uPRO) combined with computational procedures to discover cell type specific enhancers, promoters, and transcription factor networks. uPRO is composed of adaptor ligation and reverse transcription reactions, which is reduced to a one-day procedure and makes nascent RNA-seq more feasible and flexible for a widespread use. We generated genome-wide profiles of nascent transcription in human blood derived cell lines and clinical samples of ~1 ml of untreated whole blood. We integrated these data into deep learning and hierarchical network analysis to detect enhancers, promoters, and co-expression networks to define cell-type specific transcription programs. We found conservation of position but variation of expression in cell type specific enhancers and transcription start sites. Transcription factors (TFs) such as TCF-3 and OCT1 were pivotally associated with TF-enhancer-gene networks across cell types. Intriguingly, we also discovered that TFs related to cell stress and inflammation - such as SRF, ATF, CHOP, and NF-kB - are associated with inter-individual variation of leukocyte transcription in whole blood. Our integration of experimental and computational nascent RNA methods will provide an efficient strategy to identify specific transcriptional programs, both in cell-type and patient/disease-associated, with minimal sample requirements.

## Introduction

Mapping active RNA polymerase is one of the most effective strategies to detect and measure transcription qualitatively and quantitatively^1^. Transcription is a converging step of gene expression that allows specific genes to respond to specific signals. Identification of these genes, regulatory enhancers, and further analysis to clarify how they are interconnected with transcription factors will lead to the revelation of control networks that specify cell types, disease susceptibility, and cellular states. Quantification of RNA polymerase activity is crucial in breaking apart and understanding these transcriptional regulatory networks^2^.

Regions of the genome in addition to protein-coding genes are transcribed to RNA: at enhancers, upstream divergent regions, and regions downstream of mRNA poly A sites. Short unstable enhancer RNAs (eRNAs) are produced from enhancers and do not encode proteins^3^, but rather identify regulatory activity of these elements^4^. Differential regulation of enhancer-mediated transcription is involved in several diseases, such as cancers, immune-related and metabolic diseases ^5^. Measuring differential transcription of eRNA is crucial in discovering master regulatory factors in nutritional, environmental, and developmental alterations. Conventional transcriptome analysis such as RNA-seq has not been efficient enough to detect such unstable RNAs. Methods specific to nascent transcription are more efficient in detecting and measuring unstable eRNAs.

Some of the existing nascent RNA methods depend on purification of insoluble chromatin^6^ or are built upon immunoprecipitation of RNA polymerase^7–9^. The specificity of these methods currently rely on antibody specificity or the efficiency of chromatin fractionation alone. However, Nuclear Run-On (NRO) based methods use nascent RNAs elongated by polymerases with nucleotide analogs and can accurately map the polymerases as well as their start sites^10–12^. Nascent RNAs are selectively labeled by nucleotide analogs using the endogenous activity of RNA polymerase. These analogs serve as affinity purification tags, providing highly specific enrichment of the nascent RNA over other forms of RNA^11^. In addition, the direction of transcription is unambiguously identified through the directional sequencing of RNA. Therefore, NRO methods are not only useful to analyze gene expression, but also to access the activities of regulatory elements and enhancers by capturing the noncoding RNAs simultaneously^3,12,13^.

The advantages in NRO methods to identify both gene expression and regulatory activity with high sensitivity and specificity make them a powerful platform to accumulate larger scale databases. Scaling up provides a lot of advantages. It can identify context specific transcription and regulatory landscape, such as in human population or disease samples^14,15^. Comparison of genotype variations and transcriptional variation can lead to revealing complex transcription related quantitative trait loci (QTLs)^15^. Large scale analysis provides high statistical power to discover disease-specific transcription profiles from patient samples. It can also differentiate networks of transcription factors to enhancers and gene expressions. Use of unsupervised machine learning could allow novel discoveries in gene expression and regulatory element architecture, provided with a large number of training data. The advantages in NRO methods to identify both gene expression and regulatory activity with high sensitivity and specificity make them a powerful platform to accumulate large scale databases.

However, there had been practical limitations in terms of the feasibility of the method and the accessibility of the samples. The Global Run-On and Precision Run-On sequencing (GRO-seq/PRO-seq) method required multiple days of hands-on procedure and robust RNA handling skills. Additionally, Nuclear Run-On requires the isolation of nuclei from intact cells, which is often a challenge for in vivo samples^12^. We previously introduced a Chromatin Run-On method (ChRO-seq) that uses insoluble chromatin isolates^14^. We demonstrated that the chromatin fraction contains enzymatically active RNA polymerases that are suitable for nuclear run-ons. Therefore, a combination of shortened PRO-seq procedure coupled to ChRO-seq using easily accessible samples would provide a powerful strategy to explore gene expression and regulatory landscape profiles in patients of specific diseases.

Here we present an ultrafast PRO-seq procedure (uPRO) coupled to peripheral blood Chromatin Run-On (pChRO) that takes less than a day to produce an Illumina compatible sequencing library. uPRO provides nascent RNA data with a quality comparable to the existing PRO-seq method. We used uPRO to explore the transcriptional landscape of various human blood derived cell lines. In particular, we report the nascent transcriptome of a myeloid derived HAP1 cell line for the first time, which has a haploid genome specifically suited for CRISPR genome editing^16,17^. We also present an integration of deep learning and hierarchical co-expression analysis to detect and define the regulatory networks of enhancers and genes. Direct peripheral blood pChRO analysis unveiled immune and stress related transcriptional programs in inter-individual variations, and demonstrated how we could deconvolute differential fractions of leukocyte subtypes from intrinsic gene expression variations using the bulk data.

## Results

### Minimalized nascent RNA-seq procedure: uPRO

#### Global transcriptional landscape of the human haploid cell line HAP1

uPRO has simplified chemical reactions RNA adaptor ligation and template switching reverse transcription that can be completed in as short as 6.5 hours (**Fig S1A**, see **Supplementary Text**). We first applied uPRO and PRO-seq on the human haploid HAP1 cell line derived from human myeloid leukemia cells^16^, and HeLa. Haploid cell lines provide an advantage over other diploid or multiploid cell lines since only one allele of the gene or elements needs to be modified. This is a critical advantage in large scale genetic screening using genome editing technologies such as TALE or CRISPR. A HAP1 transcriptional landscape profile will serve as a useful baseline resource^18^. We also compared uPRO to existing PRO-seq data from human embryonic kidney HEK293 cells and lymphoblastoid cell lines (LCLs)^19,15^. Overall, uPRO shows transcription profiles equivalent to PRO-seq results in HAP1 and in HeLa cells (**Fig 1**). For example, uPRO and PRO-seq show highly consistent transcription profiles at TAL1 gene, a HAP1 specific gene that is often up-regulated in T cell origin leukemias (**Fig 1A**)^20^. Both the sense and antisense strand transcription patterns are efficiently captured. An adjacent gene STIL is expressed in both HAP1 and HEK293 cells, and the expression pattern between uPRO and PRO-seq in HAP1 is in high agreement.

**Fig 1.**
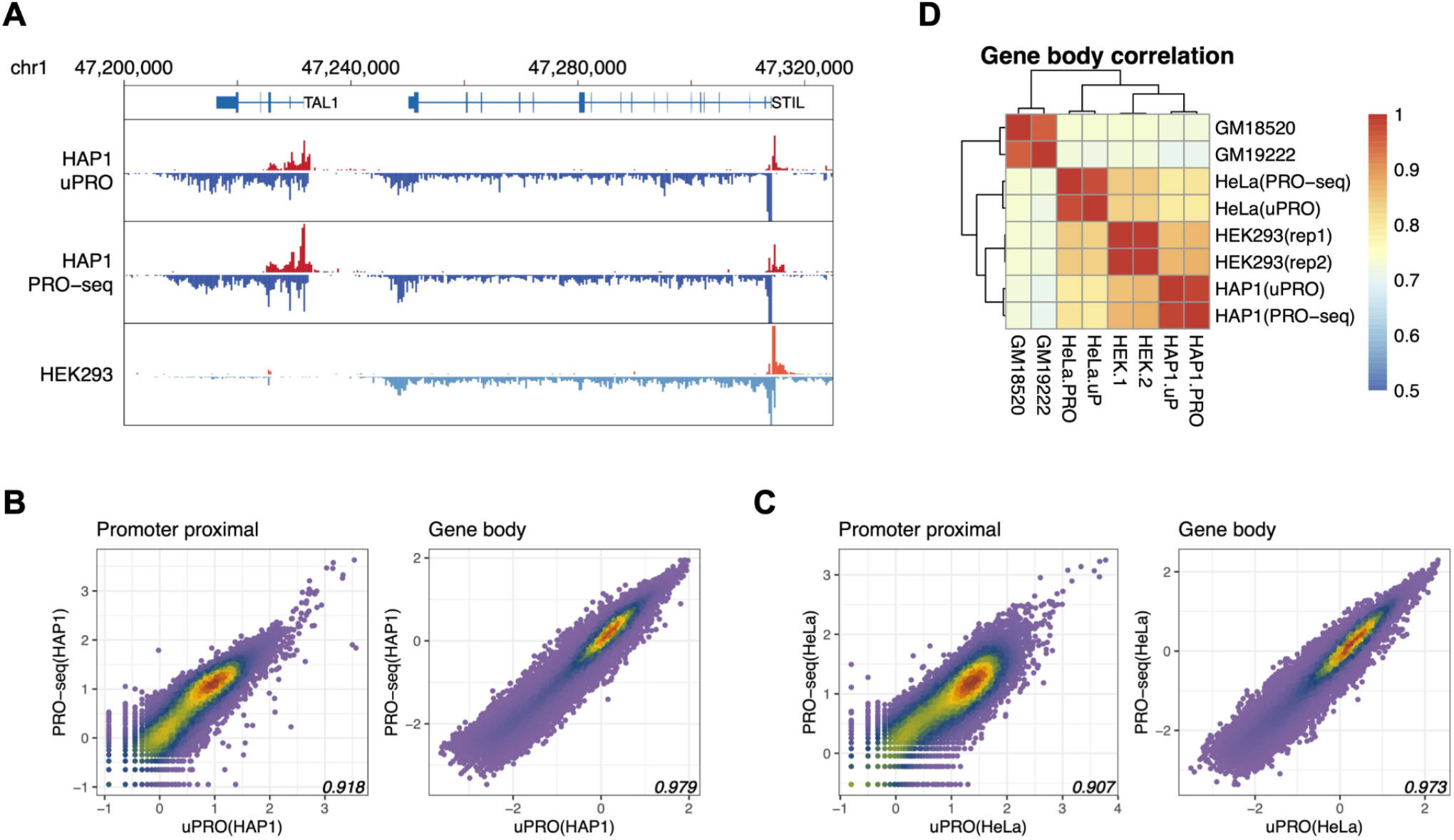
Comparison between uPRO and PRO-seq. **A.** Browser view of a representative loci showing cell type specific genes. Red: plus strand gene or PRO-seq track, blue: minus strand gene or PRO-seq track. **B.** Correlation scatterplots between uPRO and PRO-seq in the promoter proximal and gene body regions in HAP1 cells. x- and y-axes: log_10_ reads per kilobase per million mapped reads (RPKM). Pearson’s correlation coefficient indicated. **C.** Correlation scatterplots between uPRO and PRO-seq in HeLa cells. **D.** Correlation heatmaps among uPRO and PRO-seq samples from various cell types. Color scale bar represents Pearson correlation coefficient of the log_10_ RPKM reads.

To quantitatively measure uPRO’s reproducibility of previously presented PRO-seq data, we compared promoter proximal and gene body read counts. Promoter proximal regions are defined as ± 500 bp from the transcription start sites and reflect the amount of RNA polymerases that are paused. Gene body regions reflect the amount of actively transcribing RNA polymerases which represents overall gene expression levels. Correlation coefficients between regions are greater than 0.9 between uPRO and PRO-seq in promoter proximal regions and 0.97-0.98 in gene body regions in HAP1 and HeLa cells (**Fig 1B, 1C**). These correlation coefficients between uPRO and PRO-seq are slightly less than between PRO-seq replicates in HEK293 cells (**Fig S1D**), but still demonstrates that uPRO quantification is a reasonably close estimate of PRO-seq quantification, in particular on the gene bodies. When we included other blood cell derived LCLs (GM18520, GM19222) from different individuals in the hierarchical clustering analysis, uPRO and PRO-seq results cluster together within the same cell lines and the clustering isn’t affected by the method (**Fig 1D**). This demonstrates the robustness of uPRO in agreement with PRO-seq.

### Deep learning based detection of cell type specific enhancers and genes

#### Artificial neural networks detect eRNA and gene TSSs in high resolution

Both gene promoters and enhancers are capable of driving transcription at the site in a divergent bidirectional manner. This bidirectional signature has been used in strategies to conveniently identify active transcriptional enhancers. Nascent RNA-seq such as PRO-seq is particularly well suited for detecting unstable enhancer RNAs. Machine learning based methods, such as Support Vector Machines (SVMs), have been used to detect the divergent bidirectional signatures of nascent RNA transcriptions at enhancers and promoters. We developed a new deep learning based Bidirectional Transcription Scan (deepBTS) tool, which shows increased performance in terms of sensitivity, specificity, and speed (1-2 orders of magnitude) using minimal computational power.

The plus and minus strand PRO-seq data were converted into input vectors using multiple layers of binning windows and normalizations (**Fig 2A,** see **Supplementary Text**). We trained neural networks using the PRO-seq data in human lymphoblastoid cell lines (LCL) to predict enhancer and gene TSSs (**Fig S2**). LCL specific TSS references are derived from previous capped nascent RNA sequencing (PRO-cap) data. The deepBTS neural networks were iteratively trained using the Fast Artificial Neural Network library (FANN), to optimize for the highest area under the receiver-operating curve (AUROC) in the precision recall analysis of the TSS references (**Fig 2B**). We tested variable binning sizes, window ranges, and network designs to select the network with the highest AUROC (**Fig S2,** see **Supplementary Text**). Compared to the SVM based method (dREG), deepBTS performed significantly with less computational resources (average of 2.0 core-minutes compared to ~10 core-hours), while yielding comparable results in terms of sensitivity and specificity (**Fig 2B**). For example, both dREG and deepBTS identified a HAP1 cell-type specific distal enhancer cluster near the CWC22 gene locus (**Fig 2C**).

**Fig 2.**
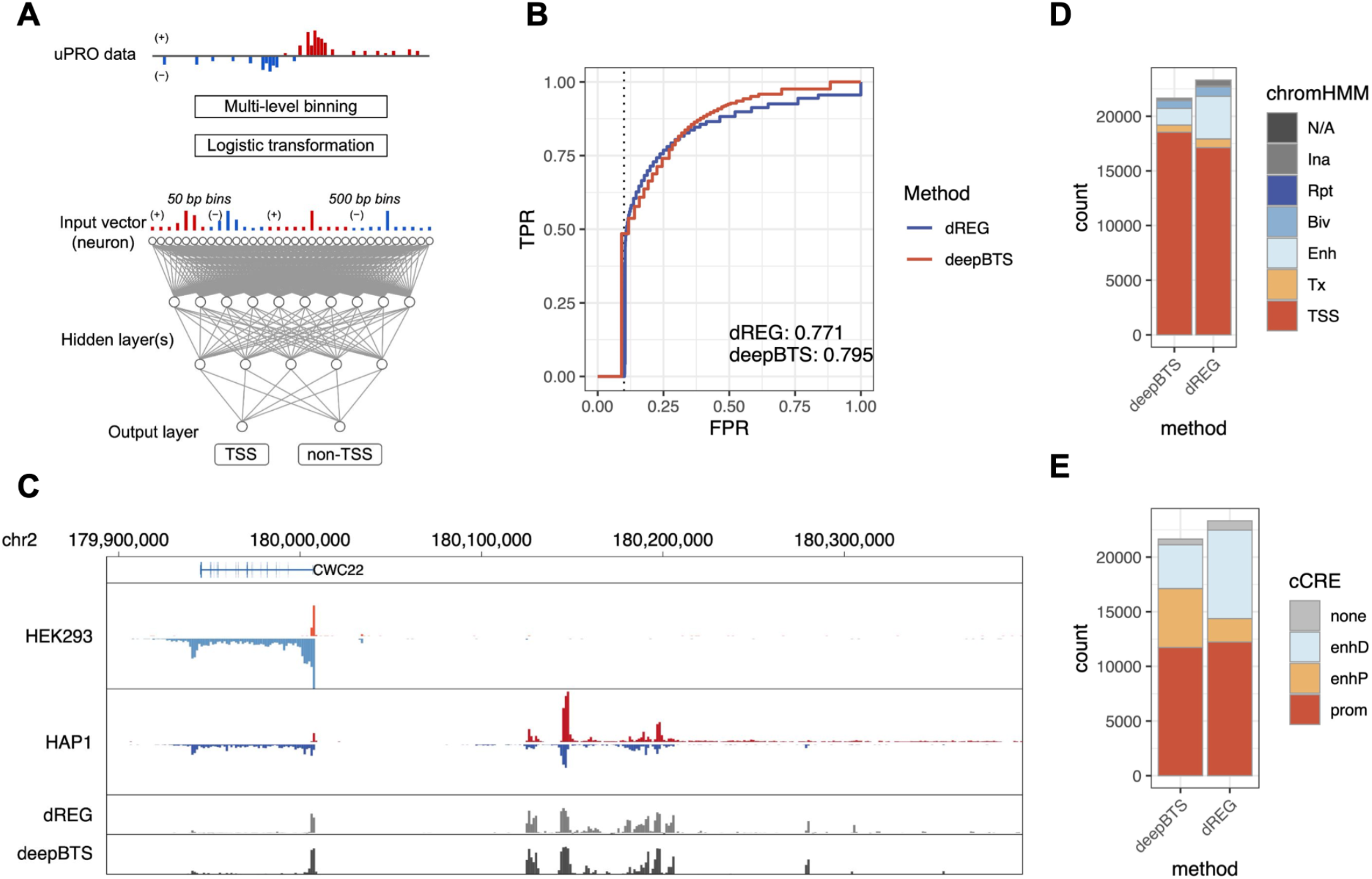
Deep learning for detecting enhancer and gene TSSs. **A.** Schematics of artificial neural network (ANN) design. **B.** Precision-recall analysis of deepBTS. AUROCs are labeled. Dotted vertical line indicates FDR = 0.1. **C.** Genome-browser view of deepBTS scores at a HAP1 specific enhancer cluster. **D.** Distribution of deepBTS TSSs with respect to chromatin domains in HEK293 cells (ChromHMM). **E.** Distribution of deepBTS TSSs with respect to ENCODE candidate cis-regulatory elements (cCREs).

Most of the bidirectional enhancer and gene TSSs identified by deepBTS are located in transcriptionally active regions of the genome. We used histone ChIP-seq data in HEK293 cells to generate chromHMM domains and assigned the chromHMM identities of the TSSs. DeepBTS identified 21,650 TSSs, of which 86% were at active TSS regions and 7% at enhancer regions, whereas dREG identified 23,317 TSSs, with 73% at TSSs and 17% at enhancers (**Fig 2D**). Small fractions (1.1% of deepBTS and 2.7% of dREG) of the TSSs were at transcriptionally inactive regions in chromHMM (inactive, repetitive, or not assigned, **Fig 2D**). While deepBTS found more chromHMM TSSs and less chromHMM enhancers than dREG, higher resolution breakdown using ENCODE candidate cis-regulatory element (cCRE) data showed that deepBTS detected more proximal enhancers (enhP, closer than 2 kb from TSSs) than dREG (**Fig 2E**). Using the cCRE data, the fraction of enhancers discovered by deepBTS and dREG are overall comparable (43% and 44% respectively). However, a larger fraction of deepBTS enhancers are proximal enhancers (57%), while dREG enhancers are less proximal (21%) and mostly distal (**Fig 2E**). This shows that DeepBTS and dREG may have distinctive performances in discovering proximal versus distal enhancers.

#### Identification of blood cell-type specific genes and regulatory elements

We applied deepBTS to 39 nascent RNA sequencing datasets (uPRO or PRO-seq) from 6 blood derived cell types (PBMC, PMNL, MDM, LCL, HAP1, THP1) and 2 outgroup cell types (HEK293, HeLa). We discovered an average of 15,625 TSSs per dataset. On average, 56% of TSSs were promoters, 19% proximal enhancers, 22% distal enhancers, and 3% did not overlap with ENCODE cCREs in each dataset (**Fig 3A, Fig S3A**). The number of TSSs detected by deepBTS increases with the library size up to 20 million reads, but reaches saturation at 20 million (**Fig 3B**). In total, 54,560 TSSs were identified, 29% were promoters, 9% proximal enhancers, 52% distal enhancers (**Table S1**). The larger proportion of distal enhancers in total count shows that distal enhancer expression is more diverse across cell types, which is consistent with the notion that distal enhancers specify cell types.

**Fig 3.**
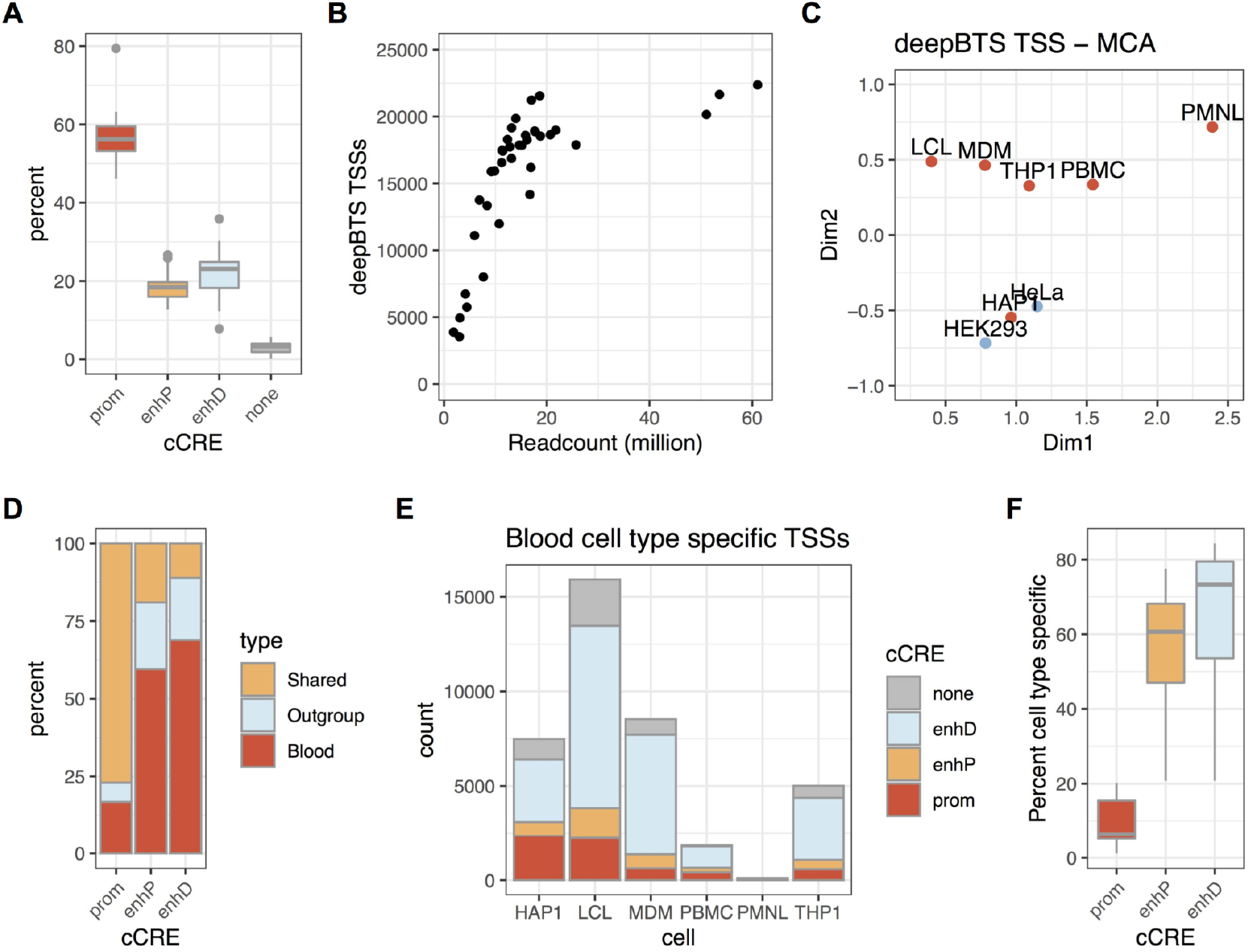
Identification of cell-type specific gene and enhancer TSSs. **A.** Proportion of cCRE classes of the deepBTS TSSs in 39 datasets. **B.** Number of deepBTS TSSs found per PRO-seq (uPRO) unique molecular read counts. **C.** Multiple correspondence analysis (MCA) of deepBTS TSS in the cell types. **D.** Cell type specificity of promoter and enhancer TSSs. **E.** Number of individual cell type specific TSSs. **F.** Fraction of individual cell type specific TSSs.

To identify cell type specific promoters and enhancers, we evaluated the overlap between the total of 54,560 TSSs in each of the 6 blood derived and 2 outgroup cell types. 14,177, 3,884, 21,672, 29,512, 20,154, 18,213, 23,513, and 18,021 TSSs were found in in PBMC, PMNL, MDM, LCL, HAP1, THP1, HEK293, and HeLa datasets, respectively. Multiple correspondence analysis of the TSS overlaps show that the blood cell types are distinguished from the outgroup cells with the exception of HAP1 cells (**Fig 3C**). Two of the monocytic-macrophage cells THP1 and MDM cluster closely, reflecting that the crude overlap of deepBTS TSS shows cell type specificity.

To identify cell type specific promoters and enhancers, we evaluated the overlap between the total of 54,560 TSSs in each of the 6 blood derived and 2 outgroup cell types. 14,177, 3,884, 21,672, 29,512, 20,154, 18,213, 23,513, and 18,021 TSSs were found in in PBMC, PMNL, MDM, LCL, HAP1, THP1, HEK293, and HeLa datasets, respectively. Multiple correspondence analysis of the TSS overlaps show that the blood cell types are distinguished from the outgroup cells with the exception of HAP1 cells (**Fig 3C**). Two of the monocytic-macrophage cells THP1 and MDM cluster closely, reflecting that the crude overlap of deepBTS TSS shows cell type specificity.

The 6 blood derived cells combined yielded 47,009 TSSs, of which 29,823 are unique to the blood derived cells. Majority (65%) of the blood cell type specific TSSs are distal enhancers (enhD; n = 19,475), while only 8.8% were gene promoters (n = 2,614). Most gene promoters are conserved across cell types (77%), but higher fractions of proximal and distal enhancers are blood cell type specific (69%, and 60% respectively; **Fig 3D**). We identified individual cell type specific TSSs (present in 2 or less of the 6 blood derived cells), and observed similar results showing the enrichment of the proximal and distal enhancers in individual cell type specific TSSs (**Fig 3E,F; Fig S3**).

#### Quantitative analysis of cell type specific enhancers and genes

Since most promoters are conserved across cell types, we speculated that cell type specific genes are regulated quantitatively at varying expression levels rather than through binary on or off control. We measured the nascent RNA expression levels at the 54,560 gene and enhancer TSSs in the 39 datasets. For intragenic or proximal enhancers overlapping or near an active gene, we quantified only the opposite strand transcription to avoid the interference from gene body or upstream antisense transcription.

The read count quantification at enhancer and promoter TSSs shows clustering of the same cell types with high correlation (**Fig 4A**). We observed the same in gene bodies (**Fig S4A**). The quantification across the gene body performed better than at the TSSs showing higher correlation coefficients between biological replicates, since they have larger regions to count reads. The gene body quantification was further enhanced by using read coverage than raw counts (**Fig S4B**). This was due to the reduction of noises, under the assumption that multiple active elongating RNA polymerases are not likely to occupy the same position by chance, and reads at the same position of the gene body are mostly duplicated reads or non-elongating RNA polymerases paused at internal enhancer TSSs. HAP1 cells showed greater similarity to HEK293 and HeLa cells, consistent with the TSS overlap MCA analysis.

**Fig 4.**
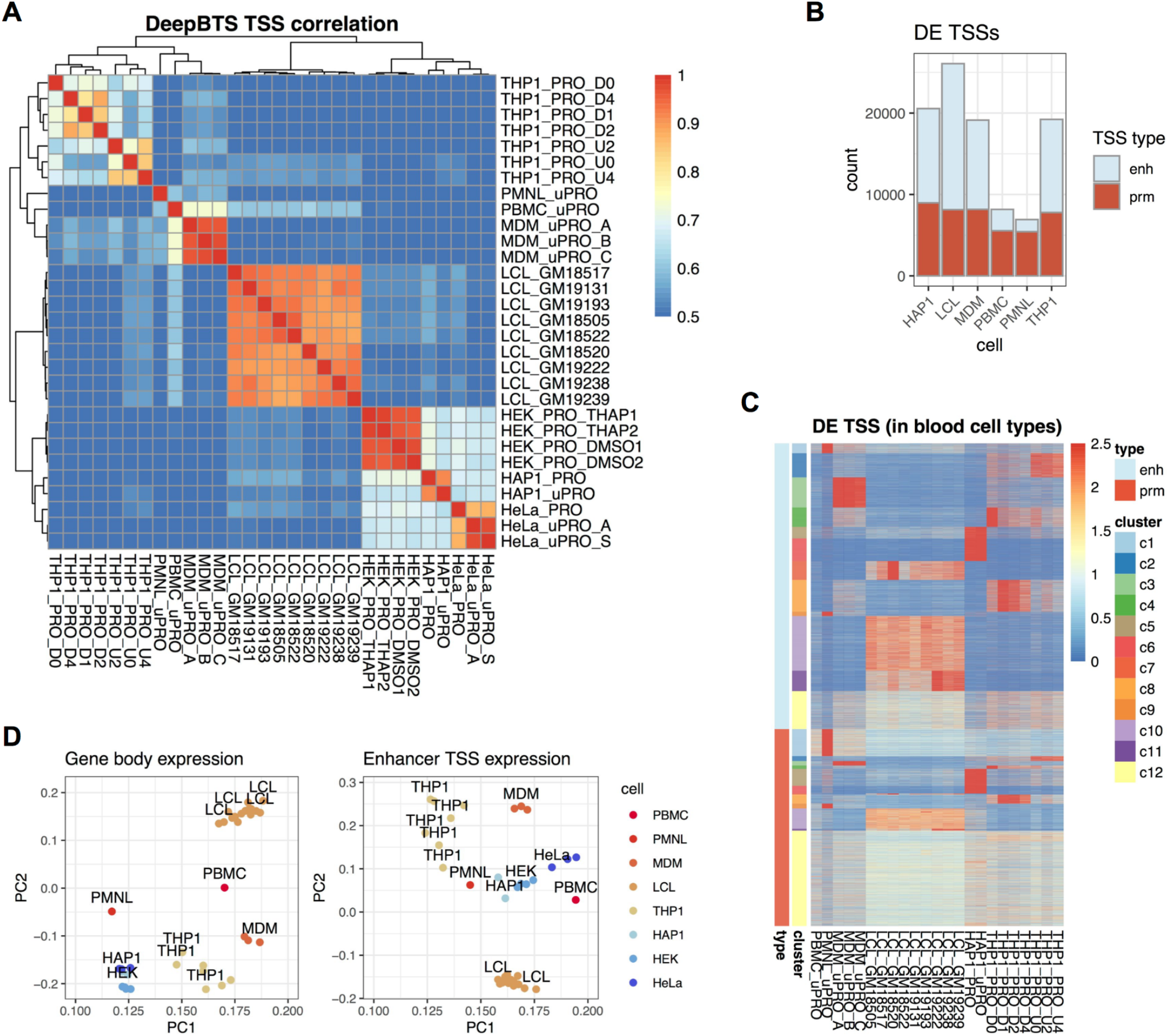
Quantitative analysis of cell type specific TSSs. **A.** Correlation coefficients of the TSS read counts across the datasets. **B.** Number of differentially expressed deepBTS TSSs. **C.** Heatmap of differentially expressed genes in blood derived cell types. **D.** Principal Component Analysis of gene and enhancer expression

We used differential expression analysis to identify cell type specific enhancers, promoters, and genes quantitatively (**Table S2**). Upon the threshold of at least 2 fold difference and false discovery rate of 0.05 using DESeq2, we found 63% of the genes are differentially expressed from the outgroup cells in at least one of the 6 blood derived cell types. At the TSSs, 71% are differentially expressed overall including distal enhancers (enhD, 59%), proximal enhancers (enhP, 66%), intragenic enhancers (enhG, 73%), and promoters (86%) . The high fraction of differentially expressed enhancer TSSs is consistent with the deepBTS cell type specific enhancer discovery. High fraction of differentially expressed promoter TSSs and gene bodies shows that cell type specific gene expression is a quantitative trait. Similarly, 60-80% of genes and TSSs are differentially expressed within the blood derived cell types (**Fig 4B, 4C**). Overall, we observed greater degree of cell type specific differential transcription at enhancer compared to promoters. For example, clusters 7, 10 and 11 contain LCL specific TSSs, mostly with enhancers that are expressed higher than average across the 6 cell types. Similarly, cluster 3 contains TSSs that are highly expressed MDMs, which is concentrated on the enhancers. Clusters 4, 8, and 2 are THP1 specific but are dependent on the differentiation and activation states (**Fig 4C**). Principal component analysis at the gene bodies, enhancer and promoter TSSs all show cell type specific clustering (**Fig 4D, Fig S4C, S4D**)

While HAP1 cells are originally leukemia derived cell lines, it showed a greater degree of similarity to the 2 outgroup cell lines HEK293 and HeLa (**Fig 4A**), presumably related to the acquisition of adherent growth phenotype. To identify HAP1 specific enhancer and genes, we compared HAP1 to HeLa or HEK293 cells. 8,565 genes are differentially expressed in HAP1 (**Fig S4E**), showing that more than 10% of the expressed genes are involved in cell type specification in HAP1 cells. 3,469 promoters (23%) are differentially expressed (1,968 up-regulated) in HAP1 cells (**Fig S4F**), indicating that a large fraction of active promoters are cell type specifically regulated. Likewise, 2,644 distal enhancers (24%) are differentially expressed (1,557 up-regulated) (**Fig S4G**) in HAP1 cells.

### Integrative analysis of cell type specific transcriptional networks

#### Hierarchical coexpression analysis of transcription factor(TF)-enhancer-gene networks

Gene coexpression network (GCN) analysis discovers genes and modules of genes that have similar expression patterns, which can identify transcriptional regulatory programs and functionally related genes. Our nascent RNA analysis can measure gene expression and the activity of cis-regulatory modules through enhancer RNA quantification. This allows the integration of GCN analysis to a deeper layer. We quantified the expression of TFs, identified binding sequences at the deepBTS enhancers and promoters (ENCODE conserved TFBS database), and mapped them to the proximity-based cis-interacting target genes to construct a framework of TF-enhancer-gene network (**Fig 5A**). We identified coexpression patterns in this framework across 29 datasets in 5 blood derived cell types. HAP1 cell was removed from this list, since it showed greater similarity to the 2 outgroup cell types.

**Fig 5.**
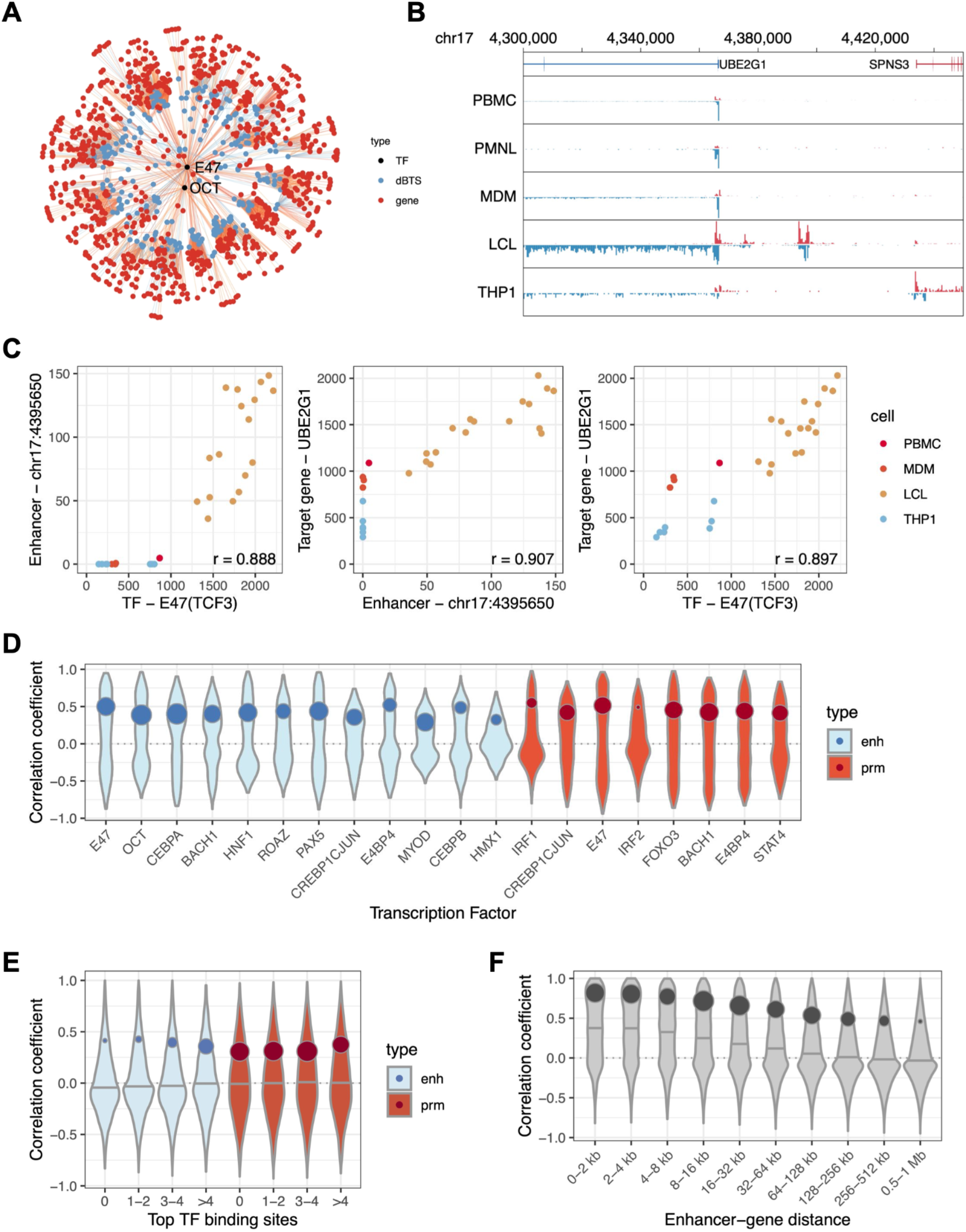
Integrative co-expression analysis of transcriptional networks. Dot colors indicate element type. Line colors indicate strength of coexpression. Only significant linkages are drawn. **A.** Example of TF-enhancer-gene coexpression network in chromosome 17. **B.** Browser example of a coexpressed deepBTS enhancer (chr17:4395650) and a gene (UBE2G1) in blood derived cells. **C.** Example of a hierarchical correlation among TF, enhancer, and target gene expression. **D.** Bimodal-violin plots of TF-TSS coexpression coefficients. Plot contour reflects the distribution; circles positions are at the upper bimodal peak, the size corresponds to the weight of the peak in a bimodal fitting. **E.** Bimodal-violin plots of coexpression between enhancers/promoters and TFs with different numbers of non-cognate binding sites (x-axis). **F.** Bimodal-violin plots of distance dependent enhancer-gene coexpression.

For example, we identified a deepBTS distal enhancer located about 30 kb upstream of the UBE2G1 gene promoter (**Fig 5B**). This enhancer contains the TF binding sequence for the TCF3 transcription factor, and its eRNA expression is highly correlated with TCF3 gene expression (**Fig 5C**, left). The eRNA expression is also highly correlated with the UBE2G1 gene expression, reflecting a cis-regulatory interaction (**Fig 5C**, middle). While UBE2G1 promoter does not contain a TCF3 binding site, UBE2G1 expression is also highly correlated with TCF3 expression (**Fig 5C**, right), suggesting TCF3 associated UBE2G1 regulation is mediated through this distal enhancer.

We further surveyed the TF-enhancer and TF-promoter coexpression patterns. From the combination of 170 ENCODE TFBS transcription factors and 55,700 enhancer or promoter TSSs, we calculated over 700 thousand TF-TSS coexpression coefficients. On average, enhancers contain 8 TF binding sites, and promoters contain 20 TF binding sites. 50 TFs showed statistically significant (fdr < 0.05) and consistent (|mean r| > 0.1) correlations to enhancer and promoter expressions (**Fig 5D**). Overall most TFs are activating TFs but we identified AHR, CP2, and ZID as repressive TFs that show negative co-expression. Even for activating TFs, the overall correlation is bimodally distributed - mixture of highly correlated TF-enhancer or TF-promoter pairs fractions and non-correlated fractions. This reflects that every TF binding site shows co-expression with the TF, and suggests the context specificity of TF function.

Overall, TF-enhancer co-expression is weaker than TF-promoter co-expression. However, some TFs such as TCF3 (E47) and OCT1 show enhancer preference. TCF3-enhancer co-expression was stronger than TCF3-promoter, possibly indicating that TCF3 is more specialized for distal enhancers. Also, motif-containing enhancers or promoters that correlated stronger with their cognate TFs tend to have arrays of other TF binding sites. Enhancers with the co-expression coefficients greater than 0.5 had an average of 13 TF binding sites, while those with lower co-expression coefficients only had 6. Conversely, enhancers containing more TF binding sites for other TFs tend to be better co-expressed with the weaker TFs (**Fig 5E**). This supports a cooperative TF binding model at enhancers and promoters.

Co-expression patterns between enhancers and target genes are also dependent on the genomic contexts. The distance between the enhancer and the target gene is expected to be one of the strongest determinants of enhancer action. We tested the correlation between nascent RNA expression levels at distal enhancers and the gene bodies of their nearest TSSs. Only the upstream intergenic DEs are tested to remove confounding effects from the gene body transcription of other genes. We observed significant correlation of expression levels between the DEs and their nearest target genes. This indicates that the DEs and their paired genes are co-regulated and that the DEs are likely regulators. We further categorized the DE-gene pairs dependent on their distances, and saw overall decay in correlation over the distance (**Fig 5F**). The decay pattern shows that an average DE-gene co-expression can span up to ~500 kb, which is consistent with other functional and chromatin conformational studies of enhancer-gene interactions.

### Direct analysis of human whole blood chromatin

#### Chromatin Run-On from peripheral blood samples (pChRO)

While cell lines are reliable sources for transcriptional profiling, the variety of established cell lines are limited. Using a large scale study will increase the analytical power in discovering inter-individual or disease associated gene expression differences. But it is not always feasible to generate cell lines (primary cells or iPSCs) from clinical subjects at a large scale. One of the most accessible clinical specimens is peripheral blood. Gene expression analysis performed directly on blood samples may provide a feasible approach in large scale studies. However, blood plasma has high-concentrations of nucleases that degrade the quality of RNA. Additionally, over-abundance of globulin mRNAs from red blood cells (RBCs) complicate RNA expression profiling experiments. Isolated leukocytes can be used for RNA expression of other transcriptional assays such as ChIP or chromatin accessibility assays, but the isolation step itself can add variability and be laborious to implement at a large scale.

Chromatin Run-On (ChRO) is an alternative Nuclear Run-On (NRO) based assay that uses precipitated chromatin isolates that contain active RNA polymerases. ChRO-seq is able to successfully generate nascent RNA data from cryopreserved archived solid tissue specimen, despite that RNA is degraded over time from these harsh conditions, because RNA polymerases can remain actively engaged. With length extension of the nascent RNAs in NRO, RNA polymerase levels and positions can be mapped even under severe RNA degradation conditions. We applied this strategy to human peripheral blood samples. Since leukocytes unlike RBCs contain a nucleus, ChRO-seq results will reflect the transcriptional landscape of peripheral leukocytes. We were able to successfully generate ChRO-seq libraries from just 1 ml of peripheral blood samples that did not undergo any special treatment but simple storage in −20°C after sampling (pChRO).

The resulting pChRO data shows high correlation in gene body read counts (0.97 - 0.98) within technical three replicates (**Fig S5A**). We also compared pChRO data from a different individual and observed slightly less but correlated gene body levels (0.95 - 0.96). The pChRO profile is reproducible between different individuals and within cell types. For example, the expression of IKZF1 gene, a known regulator of lymphocyte differentiation, is consistently high in the pChRO data from both individuals but is not expressed in HeLa cells (**Fig S5B**). On the other hand, the FIGNL1 gene is only expressed in HeLa cells but not in either of the pChRO data from the two individual samples. When we tested the gene body correlation between the pChRO replicates and individuals against HeLa and LCL samples, we saw clear clustering of pChRO data among the pChRO replicates (**Fig S5C**). The pChRO data reflecting peripheral leukocytes correlate relatively better with LCLs, B cell derived cell lines, than HeLa cells indicating that the cell type lineage is still preserved in immortalized LCLs.

#### Leukocyte decomposition between Peripheral Blood Mononuclear Cells (PBMCs) and Polymorphonuclear leukocytes (PMNLs)

One of the challenges in identifying differentially expressed genes and regulatory elements from primary cells such as in whole blood is cellular heterogeneity. If there are factors that influence cell type compositions, this will lead to false positive identification of differentially expressed genes. Peripheral leukocytes are composed of PBMCs that include B, T lymphocytes, and monocytes; PMNLs are considered as the granulocyte population composed of neutrophils, eosinophils, and basophils. The nuclear morphology and gene expression profiles are known to be different between the two, but there has been no systematic comparative transcription analysis. In particular, the gene expression profiles of PMNLs are not as extensively studied as PBMCs.

To address this, we used the uPRO data from freshly isolated PBMCs and PMNLs. The majority of the transcription profile appears to be similar between PBMC, PMNL, and whole leukocyte. However, we found genes that are differentially and almost exclusively expressed in one cell type versus the other (**Fig 6A**). For example, the ALPL gene is exclusively expressed in PMNLs but not in PBMCs. Therefore, since the level of ALPL expression is specific to PMNL cells, its expression level should also be reflected from the amount PMNL cells in the whole blood.

**Fig 6.**
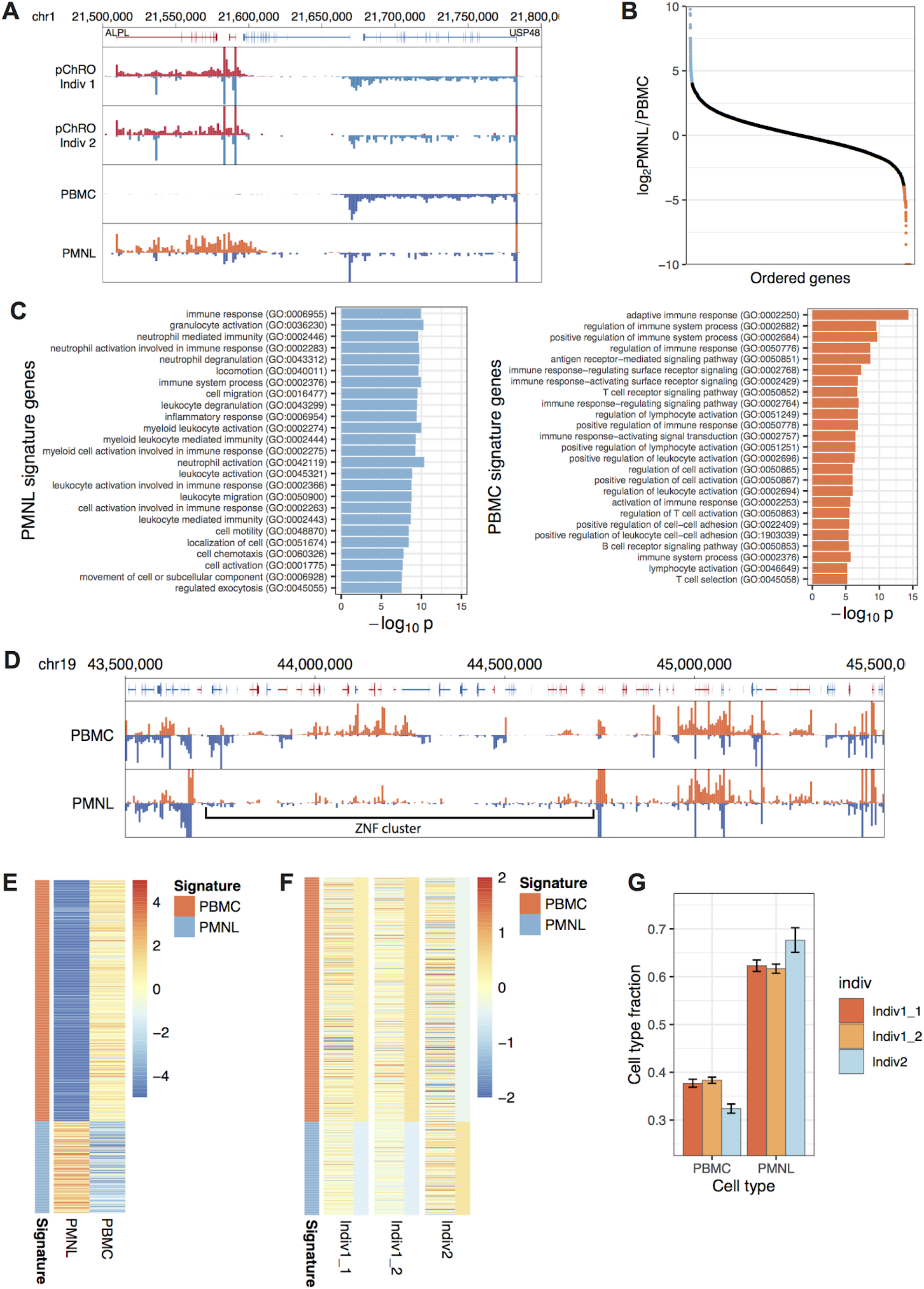
Decomposition of heterogeneous blood cell types between PBMC and PMNL. **A.** Browser view of an example exclusively expressed site. Red: plus strand gene or PRO-seq track, blue: minus strand gene or PRO-seq track. **B.** PMNL/PBMMC ratio of all genes. Blue: PMNL signature genes, red: PBMC signature genes. Signature gene cut-off: greater than 16 fold difference. **C.** Gene ontology (GO) catagories of PMNL and PBMC signature genes. Top 25 are shown **D.** Browser view of the ZNF cluster at chr19q13.31. Red: plus strand gene or PRO-seq track, blue: minus strand gene or PRO-seq track. **E.** Heatmap of signature gene expression in PMNL and PBMC uPRO gene body normalized to whole leukocyte pChRO levels. Color scale bar: log_2_ fold difference. **F.** Heatmap of signature gene expression in pChRO individual samples. Thin ribbons on the right represent the average fold difference of the signature gene group in the corresponding individual. Color scale bar: log_2_ fold difference. **G.** Estimated cell type fractions calculated from signature gene expression levels. Error bars: standard error of the mean.

On the other hand, a nearby gene USP48 is more highly expressed in PBMCs compared to PMNLs (**Fig 6A**). Quantitative assessment of these exclusively expressed signature genes should allow us to precisely estimate the PBMC and PMNL ratio in the whole blood.

To identify PMNL or PBMC exclusive genes, we calculated the ratio between PMNL and PBMC normalized gene body read counts. We identified 157 PMNL and 429 PBMC signature genes that have at least 16 fold expression differences (**Fig 6B**). Gene ontology analysis of these signature genes were consistent with the expectation. For example, PMNL signature genes are enriched with granulocyte activation (GO:0036230), neutrophil involved pathways (GO:0002446, 0002283, 0043312, 0042119), and cell motility/migration (GO:0040011, 0048870, 0050900) (**Fig 6C**). These pathways are consistent with the function of granulocytes and neutrophil innate immune responses. On the other hand, PBMC signature genes are enriched with adaptive immune response (GO:0002250), receptor mediated immune signaling (GO:0050851, 0002768, 0002429), T cell pathways (GO:0050852, 0050853, 0045058), and B cell pathway (GO:0050853) (**Fig 6C**). This is consistent with the fact that PBMCs are mostly composed of T cells and B cells.

Interestingly, we found that many of the top PBMC signature genes are among the ZNF subfamily genes. These genes are clustered on chr19 q13.31 ~1 megabase region (chr19: 43,700,000 - 44,700,000). We found that the whole 1 megabase region is repressed in PMNLs but is expressed in PBMCs (**Fig 6D**). This variation in transcription could potentially lead to a mis-interpretation that a factor affecting the PBMC and PMNL ratio may appear to influence the large range repression of this ZNF cluster. This case illustrates the importance of deconvoluting cell type heterogeneity in peripheral blood gene expression profiling.

#### Decomposition of leukocyte subtype fractions from pChRO and reference PBMC/PMNL data

For exclusively expressed signature genes, the ratio between normalized pChRO data and the signature cell type will reflect the relative cell fraction in the whole blood leukocyte population. Since PMNLs normally compose 65% of the leukocyte population, we should expect to see the pChRO data recapitulate the PMNL profile more than the PBMC profile. However, PBMC profiles are closer to the whole blood pChRO profile and log_2_ fold difference is closer to 0 after normalizing to the pChRO data (**Fig 6E**). This indicates that PBMCs are transcriptionally much more active than PMNLs; Even though PMNLs compose greater fraction of the cells, PBMCs override the transcriptional profiles. We calculated the relative transcriptional activities and PBMCs are on average ~ 4.1 fold transcriptionally more active than PMNLs. Regardless, the expression difference between the two cell types is pronouncedly distinct in the signature genes (**Fig 6E**).

To calculate the PBMC and PMNL compositions, we calculated the average expression level of signature genes relative to the reference pChRO data. This average is proportional to the relative fraction of the signature cell type. From the pChRO profiles of the individuals 1 and 2, Individual 2 appears to have a slightly higher overall expression level of PMNL signature genes than Individual 1 (**Fig 6F**). We were able to calculate the cell subtype fractions from these signature gene averages (**Fig 6G**). Individual 2 has marginally but significantly higher levels of PMNL fractions while the two technical replicates of Individual 1 showed overlapping error margins. We extended this signature deconvolution approach, and used existing mRNA microarray data in PBMC subtypes^26^. Further subclassification of PBMC subtypes showed similar enrichment of granulocyte (GRAN) in individual 2 (**Fig S6**, see **Supplementary Text**).

#### Gene and enhancer transcriptions are variable between individuals

We further analyzed 12 peripheral blood Chromatin Run-On (pChRO-seq) datasets, generated from 0.5 - 1.0 ml frozen blood samples in 10 de-identified individuals. Gene expression in immune cells such as cytokine response is an important factor in host-virus interaction. Our PRO-seq data show expression levels of immune-response related genes from human peripheral blood leukocytes. Out of 3,299 “immune” related genes classified by Gene Ontology, we found 990 differentially expressed genes in at least 1 individual (false discovery rate < 0.05, DESeq). Typically, PRO-seq levels are two or more fold higher in one or more individuals in these differentially expressed genes (**Fig 7A**). These expression patterns form clusters of immune related genes and clusters of individuals (**Fig 7B)**. The co-clustering of biologically replicated data from one individual (P1, P1n, P1nh) shows the reproducibility of PRO-seq in peripheral leukocytes (pChRO). The gene expression patterns suggest that individuals can be classified into groups with different immunity patterns if PRO-seq in peripheral leukocytes were conducted on a larger scale.

**Fig 7.**
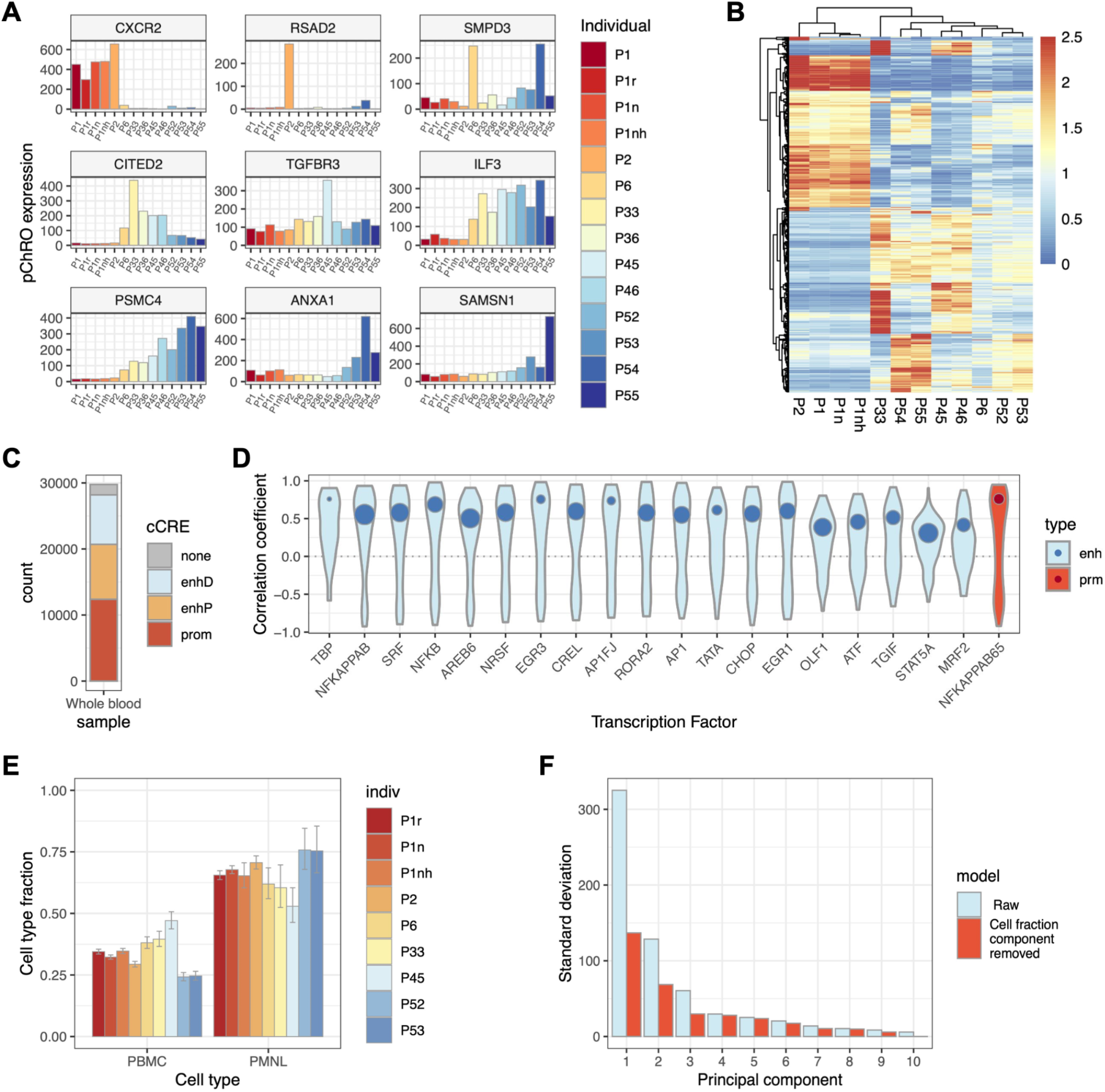
Transcriptome-wide profiles of leukocyte gene expression using uPRO. **A.** Expression levels of the most differentially expressed immune related genes in each individual. Expression levels were normalized using DESeq2, and the most differentially expressed genes were selected based on their p-values of the DESeq analysis. **B.** Heatmap of uPRO expression levels of the immune related genes and hierarchical clustering of the genes and individuals.Y-axis shows the 3,299 immune related genes clustered. Colormaps reflect the z-score normalized expression levels within the same genes across different individuals. **C.** DeepBTS TSS counts in each ENCODE cCRE category. **D.** 20 top TFs with the highest TF-enhancer or promoter coexpression coefficients. **E.** Cell type deconvolution of the pChRO dataset. **F.** Principal component analysis of variance before (Raw) and after removing the cell type fraction component.

To further dissect peripheral blood cell type specific transcriptional programs, we applied our streamlined integrative nascent RNA analysis procedure to the 12 pChRO samples. DeepBTS identified 29,765 peripheral blood enhancer and gene TSSs from the combined pChRO dataset with 36.8 million unique reads, taking 2 min 52 seconds. 90% (n = 26,843) of the peripheral blood TSSs were shared with the blood derived cell data. Of the deepBTS TSSs, 42% were ENCODE cCRE promoters, 28% and 25% were proximal and distal enhancers respectively, and only 5.3% did not map to cCREs (**Fig 7C**), demonstrating a robust level of discovery rates. From the deepBTS TSSs, we defined enhancer and promoter regions for quantitative analysis, and used DESeq to normalize TSS and gene transcription quantifications. The hierarchical TF network analysis reveals the top TFs involved in inflammatory processes such as subunits of NK-kB, and cell stress response factors such as SRF, CHOP, and ATF (**Fig 7D**). Cell type deconvolution analysis shows differential white blood cell composition in each individual sample (**Fig 7E**), and a significant fraction of the expression variation remains after removing the principal component to correct for the cell type fractions (**Fig 7F**). Further analysis normalizing the cellular heterogeneity from the clinical differential cell count data would distinguish the intrinsic variations in the transcriptional programs across the individuals. These results demonstrate that the integrated nascent RNA analysis approach can be directly applied to raw blood samples, and can reveal critical transcription programs responsible for phenotypic variations in peripheral leukocytes.

## Discussion

Nascent RNA sequencing is a powerful method that can map and measure gene expression and the activities of the transcriptional regulatory elements at the same time. In this study, we present the uPRO method, coupled to an integrated pipeline using deep learning and co-expression networks to measure gene expression, enhancer activity, and identify associated transcriptional programs. Also, peripheral blood ChRO-seq is expected to lower the barrier and facilitate the production of nascent RNA data in larger scale clinical samples.

Methodologically, we provide a new faster nascent RNA method with minimal reaction steps yielding equivalent results with existing methods. We applied deep learning in identifying bidirectional peaks of enhancers and promoters. The hierarchical coexpression analysis identifies transcriptional programs, above differentially expressed genes, and detects functional association of TFs and enhancers through cis- and trans-regulatory linkages. Direct whole blood pChRO expands sample feasibility, and we suggested a way to infer cell heterogeneity variation from gene expression profiles

Applying these methods in cell lines and human samples lead to a number of new biological findings. We provide the first transcriptional landscape of the human haploid cell line HAP1, an optimal model for genome editing. Cell types specific enhancers and promoters are differentially expressed quantitatively rather than as on/off switches. In blood derived cell lines, we found TCF3 and OCT1 as the TFs in the cell type specific transcriptional programs. We analyzed properties of enhancers that are more co-expressed with TFs and genes, such as cooperativity and cis-range interactions. In the direct analysis of peripheral blood cells, we provided the transcription landscape of polymorphonuclear leukocytes (PMNL) for the first time, identified PMNL signature gene expression, and reported the domain repression of the ZNF cluster. We also observed expression variation in peripheral leukocytes across individuals, and found that immune and stress related transcriptional programs are associated with peripheral leukocyte variation.

The quality of uPRO data is comparable to conventional PRO-seq, but uPRO can be processed faster. Removal and shortening of many critical enzymatic steps allowed more efficient use of library preparation time. High correlation of the gene body read counts between uPRO and PRO-seq reassures that uPRO is equivalent to PRO-seq for gene expression analyses (**Fig 1D**). Correlations in the promoter proximal read counts were slightly lower (**Fig 1B, C**), presumably due to potential biases in collecting read counts from a narrowed region near the 5′ end of the gene. One experimental concern has been the use of template switch reverse transcription, since it has been reported that template switch is more efficient on 5′ capped RNA ends than other forms of 5′ ends^27,28^. However, the difference of the promoter reads between uPRO and PRO-seq is similar to or smaller than the correlation between PRO-seq and PRO-cap^29^. Since both PRO-seq and PRO-cap use 5′ RNA adaptor ligation, we saw no systematic evidence that template switching at 5′ capped ends is more biased in uPRO.

The use of artificial neural networks in deepBTS was highly efficient in detecting bidirectional transcription signatures at enhancers and promoters. It performed similarly as the existing method dREG, but the speed was faster by orders of magnitude. Identified regions from individual cell types mostly overlapped with known ENCODE cCREs, indicating that deepBTS was robust, and that the positions of cell-type specific enhancers and promoters are conserved. DeepBTS showed the potential to detect proximal enhancers more than dREG, while dREG detected more distal enhancers (**Fig 2E**). We speculate that this difference is from the sensitivity of detecting lowly expressed eRNAs under different transcriptional contexts, as we saw that the increased sensitivity was accompanied by increased false positive rates in both deepBTS and dREG. We also tested that deepBTS can be used to infer other chromatin structures such as DNase hypersensitivity sites with AUROCs higher than 0.7, and can be modified for a higher resolution detection of TSSs.

Gene coexpression network (GCN) analysis has been a widely used method in discovering regulatory modules. GCN is a correlative analysis that may have limitations in discovering causal effects and identifying primary regulators by itself. In particular, multicollinearity has been a fundamental problem in resolving the causal regulatory linkage, when a candidate factor is correlated with other genes. In a meta-analysis of highly divergent cell types, multicollinearity can lead to difficulty in resolving regulatory modules from overriding large cell type specific clusters. Our hierarchical integration of cis-regulatory non-coding enhancers, their TF binding sites, and cognate TF expression allowed resolving regulatory transcriptional programs. This enhanced co-expression network is augmented by functional evidence of TF and enhancers, providing more information than simple differentially expressed gene clusters. We also experienced that coexpression networks may be less biased by multicollinearity when performed across more similar cell types. For example, we excluded HEK293, HeLa, and HAP1 that were more distinct (**Fig 4A**) for the hierarchical coexpression network analysis, and observed higher p-values of co-expression in cognate TF-enhancer pairs than including HAP1. We suspect removing outgroup samples helped resolving regulatory co-expression from multicollinear correlation. In that sense, direct pChRO analysis in even a larger scale than we performed in this study would provide an extremely useful dataset to discover transcriptional programs with high confidence with this approach.

We further explored features of distal enhancers that are associated with stronger co-expression with cognate TFs and target promoter. Enhancers coexpression with their cognate TF or target gene is probably context dependent, since coexpression is bimodally distributed. We only tested a few possibilities, cooperativity and cis-interaction distance in this study. Testing context specific effects on co-expression would be a target of further study with a larger scale analysis. In particular, the correlations between enhancers and genes were negatively associated with the distance as expected up to ~500 kb (**Fig 4F**). The degree of distance dependence can also be context dependent. For example, distance dependency can vary chromatin state or TF binding on or between enhancer and genes, such as insulating effect by CTCF. It is also possible to calculate pairwise co-expression against a specific cell type, and derive cell-type specific cis-effect ranges. For example, polyploid cancer cell lines such as HeLa cells^30^ may have more rapid distance-decay than haploid HAP1 cells, somewhat consistent with a study reporting fewer short range interactions (< 1 Mb) in Arabidopsis tetraploid cells compared to diploid cells^31^.

Peripheral blood is one of most accessible human specimens, and can be a valuable resource to obtain large scale data. However, there are often limitations due to the presence of plasma nucleases and the requirement to isolate leukocyte from massively abundant red blood cells. Our pChRO procedure is well optimized for this purpose requiring no sample pretreatment and only very simple preparation steps. In turn, we successfully generated whole blood leukocytes as well as leukocyte subpopulation data. In particular, polymorphonuclear leukocytes (PMNLs) including neutrophils are known to be transcriptionally inert, and we have quantitatively analyzed the global transcriptional activity. We also identified PMNL specific expressed genes as well as the large domain repression as seen in the ZNF cluster. The large domain repression may be related to the segmentation of the nuclei in neutrophils and would provide further insight in linking gross nuclear morphology with molecular events taking place at these domains^32,33^.

The pChRO data showed consistency between technical replicates, and identified differentially expressed genes between different individuals. We also demonstrated that cell subtype fractions can be calculated and provide important check-points in discovering true differentially expressed genes. Furthermore, we applied the integrated analysis using deepBTS and hierarchical coexpression network to define peripheral leukocyte enhancers and identify significant transcriptional programs associated with inter-individual variation. As expected, immune related TFs such as NFkB subunits show strong association. But the significance of cell stress response related TFs provides a novel insight that different leukocyte stress levels may play a role in phenotypic variations in human individuals. While our analysis had been limited due to the complete de-identification of the samples, associated clinical data would greatly expand the use of our analysis on disease state or susceptibility. Therefore, in addition to identifying differential gene expressions, mapping regulatory elements, and identifying transcriptional programs, these peripheral leukocyte pChRO data may provide further information on personalized health conditions, and open up a new revenue in the studies of genome-wide transcription.

## Acknowledgements

We thank the members of the Kwak lab and John Lis lab in Cornell University for the scientific and technical discussions and reagent sharing. This study was supported by NIH 1R35GM142979 and discretionary funds (Cornell University) to HK.

## Author Contributions

This study was conceived by HK. SSYK, and EK performed the experiments; EK, AKK and HK performed the computational analyses.

## Data depository

GSE103719, GSE110638.

Source codes for data processing and figure generation can be found at https://github.com/users/kwaklab-cornell/uPRO

## Materials and Methods

### Materials

HAP1 cells were obtained from Horizon Discovery, maintained in IMDM media with 10% FBS and 1% penicillin-streptomycin. Only cells less than passage 10, mostly between 4-6 were used. Lymphoblastoid cell lines were obtained from the Coriell cell repository. THP1, HEK293, and HeLa cells were obtained from ATCC, and maintained following the manufacturer’s instructions. PMBCs and PMNLs were directly isolated from fresh human whole blood using Lympholyte cell separation media, following the manufacturer’s instructions. Monocyte Derived Macrophage (MDMs) were isolated from deidentified donors and maintained as described previously [reference Parkmin].

### Chromatin preparation from human whole blood

1 ml of frozen blood sample is thawed and lysed in 10 ml of the NUN buffer (0.3M NaCl, 1M Urea, 1% NP-40, 20mM HEPES, pH 7.5, 7.5mM MgCl2, 0.2mM EDTA, 1x protease inhibitor cocktail, 1 mM DTT, 4 u/ml RNase inhibitor) with gentle mixing. Blood chromatin is pelleted by centrifugation at 15,000 g for 20 min, 4°C, and resuspended in Wash buffer (50 mM Tris pH 7.5). After brief centrifugation, the pellet is washed once more in Buffer D (50 mM Tris-HCl, pH 8.0, 25% glycerol, 5 mM Mg Acetate, 0.1 mM EDTA, 5 mM DTT), then homogenized using short sonication cycles in 50 μl of Buffer D.

### Isolation of Peripheral Blood Mononuclear Cells (PBMC) and Polymorphonuclear Leukocytes (PMNL)

5-10 ml of peripheral whole blood is sampled from brachial veins. A final concentration of 1.5 mM EDTA is added to the whole blood to prevent clotting. To isolate PBMC and PMNL from the peripheral blood, 3 ml of Poly Cell Separation Media, 4 ml of Human Cell Separation Media, and 5 ml of blood are carefully layered consecutively with minimal mixing. Separation occurs with a centrifugation of 450-500 g for 30-35 minutes at 18-22°C. 3 interface layers should be visible. The top interface is PBMC and the middle interface including the layer right below is PMN. 5ml of each PBMC and PMN layers are collected in 1.7 ml microcentrifuge tubes. PBMC and PMN are pelleted by centrifugation at 4,000 for 4 min, 4°C, and washed in PBS. Centrifugation and wash is repeated once more. After brief centrifugation, the pellet is washed in Buffer D (50 mM Tris-HCl, pH 8.0, 25% glycerol, 5 mM Mg Acetate, 0.1 mM EDTA, 5 mM DTT), then resuspended in 50 μl of Buffer D.

### uPRO library preparation

Chromatin or cells were incubated in the nuclear run-on reaction condition (5 mM Tris-HCl pH 8.0, 2.5 mM MgCl_2_, 0.5 mM DTT, 150 mM KCl, 0.5% Sarkosyl, 0.4 units / μl of RNase inhibitor) with biotin-NTPs and rNTPs supplied (18.75 μM rATP, 18.75 μM rGTP, 1.875 μM biotin-11-CTP, 1.875 μM biotin-11-UTP for uPRO; 18.75 μM rATP, 18.75 μM rGTP, 18.75 μM rUTP, 0.75 μM CTP, 7.5 μM biotin-11-CTP for pChRO) for 5 min at 37°C. Run-On RNA was extracted using TRIzol, and fragmented under 0.2 N NaOH for 15 min on ice. Fragmented RNA was neutralized, and buffer exchanged by passing through P-30 columns (Biorad). 3’ RNA adaptor (/ 5Phos/NNNNNNNNGAUCGUCGGACUGUAGAACUCUGAAC/3InvdT/) is ligated at 5 μM concentration for 1 hours at room temperature using T4 RNA ligase (NEB), followed by 2 consecutive streptavidin bead bindings and extractions. Extracted RNA is converted to cDNA using template switch reverse transcription with 1 μM RP1-short RT primer (GTTCAGAGTTCTACAGTCCGA), 3.75 μM RTP-Template Switch Oligo (GCCTTGGCACCCGAGAATTCCArGrGrG), 1x Template Switch Enzyme and Buffer (NEB) at 42°C for 30 min. After a SPRI bead clean-up, the cDNA is PCR amplified up to 20 cycles using primers compatible with Illumina Small RNA sequencing. The whole procedure takes ~6 hours with ~3.5 hours of hands-on time.

### Sequencing data processing

First 8 bases of each sequence read in the uPRO fastq file are clipped and tagged separately as the unique molecular identifier (UMI) to the read index. The UMI tagged fastq data is aligned to the human hg38 genome using the STAR aligner. The aligned result is processed to bam format sorted by read position and reads with identical UMI at the same positions are collapsed into one read. From the UMI collapsed bam file, genome wide read count coverage files (bedgraph) were generated for both plus and minus strands. We used dREG (https://github.com/Danko-Lab/dREG)^13^ to map transcriptional regulatory elements for comparison to deepBTS results. Read counts on genes (gene body and promoters), and de novo annotated enhancers from the plus and minus strand bedgraph files using de novo gene annotations were made using the BEDtools suite (https://bedtools.readthedocs.io/en/latest/content/bedtools-suite.html). The read counts are divided by the size factor defined as the median of the ratio to the geometric mean across samples, which is the same method used in DESeq^34^.

### Deep Bidirectional Transcription Scan (Artificial Neural Network)

Genome-wide uPRO/PRO-seq read count data (bedgraph) on both plus and minus strands are binned using multiple binning sizes, and converted to input evaluation vectors for all transcribed regions (see Supplementary Text for bin and step sizes tested). Transcribed regions are defined using a heuristic criteria of at least 3 reads in any 100 bp region within 1 kb on either plus or minus strand, and at least 1 read on each of the plus and minus strand within 1 kb region. The read count input vectors (v) are normalized using a pseudo-logistic function 1 / (1 + e^-a(v- b)^), where a = 1/b * log(1/y - 1), b = x * max(r), x = 0.05, y = 0.01. Input vectors are evaluated by a pre-trained neural network using the Fast Artificial Neural Network (FANN) library. Training sets were generated using LCL PRO-seq data as the input and TSS reference from LCL PRO-cap data as the labels. Training was performed with consecutive resilient backpropagation (RPROP) and standard backpropagation (SBP) algorithms. Testing results are reported as a bedgraph file of genomic coordinates and scores between the range of −1 to 1. The genomic coordinates with scores greater than threshold are combined as deepBTS regions. Receiver-operating curves (ROC) were generated using various thresholds, and assessing the fraction of overlapping regions to the PRO-cap defined TSS reference within 200 bop as the gold standard to derive true positive rates and false positive rates. For the ROC evaluation, we used 2 testing datasets (GM19238, GM19239) that were excluded from the training sets. DeepBTS performance was tested under a UNIX compatible system with Intel Xeon CPU at 2.40GHz clock speed using single core. The deepBTS program is available at a public source code repository (github.com/kwaklab-cornell/deepBTS).

### Cell type specific enhancer and differential expression analysis

To define regions to quantify deepBTS defined enhancer and promoter activity, we considered the strand specificity and gene body overlap. We first defined active gene bodies as refFlat annotated protein coding genes that overlap with deepBTS regions at their promoters. Then we categorized deepBTS regions as distal enhancers (>2 kb upstream of gene TSS or >10 kb downstream of gene poly(A) site), proximal enhancers (200 bp - 2 kb from gene TSS), promoters (< 200 bp from gene TSS), and intragenic enhancers. To avoid the interference from gene body transcription, intragenic enhancers are quantified only using the strand opposite of the gene (antisense). For the proximal enhancers, we quantified only the sense strand reads to avoid the interference from upstream antisense transcription from the gene promoter. Promoters transcription is quantified separated as sense and upstream antisense. Distal enhancers are quantified from both strands. Raw read count tables generated on reference gene annotations or on deepBTS identified regions. The raw read count tables were processed through DESeq2^34^ and selected differentially expressed genes or dBTSs with FDR < 0.05.

### Hierarchical coexpression network analysis

To construct the hierarchical coexpression network of TF-enhancer-genes, we used HMR conserved transcription factor binding sites (TFBS) data, and identified TFBS located at deepBTS enhancers and promoters. We also linked enhancer-gene TSS pairs within 1 M base distance. Normalized uPRO/PRO-seq gene body/deepBTS region read counts in multiple datasets were used to generate TF expression, enhancer/ promoter expression, and gene expression matrices. Correlation coefficients across cell types were calculated for each pair of TF-enhancer/promoter combinations and enhancer-gene combinations. P-values were calculated from each combination using the significance of the product moment of Pearson’s correlation test. To evaluate the strength of the association between TFs and its target regulatory elements, we tested the significance of the distribution of the coexpression coefficients at false discovery rate < 0.5. To visualize the TF effect, we made a bimodal gaussian model of the distribution of the coexpression coefficients of each TF to all of its targets, and calculated the value and the weight of the fitted centers, and plotted as a circle plot overlaying the violin plot of the distribution. The R scripts used in this analysis is available at the public source repository (github.com/kwaklab-cornell/uPRO)

### Cell type signature gene selection and decomposition analysis

Gene body uPRO data from PBMC and PMNL are normalized to reads per million mapped reads per kilobase (RPKM). PBMC and PMNL signature genes were selected by log_2_ ratio between the two, using ±4 (16 fold difference) as a cutoff. Gene ontology analysis of the signature genes were performed using the PANTHER gene ontology analysis^35^. Cell type fractions were calculated from the relative expression levels of each signature gene subset relative to the reference data, assuming reference peripheral leukocyte compositions of normal healthy adult individual: PBMC = 0.35, PMNL = 0.65, B = 0.07, T = 0.21, and CD4 = 0.06^25^.

