## Supplementary Information for "Integrative nascent RNA methods to reveal cell-type specific transcription programs in peripheral blood and its derivative cells"

**Supplementary Text**

***Comparison between PRO-seq and uPRO procedures.*** Compared to PRO-seq, uPRO requires less RNA chemistry and handling steps (**Fig S1A**)[12]. In PRO-seq, Nuclear Run-On (NRO) is performed on isolated nuclei. In uPRO, the NRO reaction is performed directly on washed cells or resuspended chromatin isolates. After the NRO reaction, the biotin-labeled nascent RNA is fragmented and the buffer is exchanged to remove excess biotin-NTPs and salts. In uPRO, 3' RNA adaptor ligation takes place for 1 hour before biotin-RNA enrichment as opposed to the 6 hour - overnight ligation after biotin-RNA enrichment in PRO-seq. This change greatly shortens the amount of time spent. 2 consecutive streptavidin bead binding then takes place and an extraction is performed to enrich biotin-labeled nascent RNA. Instead of 3 streptavidin affinity purifications throughout the procedure, we found that 2 consecutive affinity purifications were sufficient to remove potential adaptor dimers and unlabeled endogenous RNAs. In PRO-seq, RNA extraction from the beads includes multiple ethanol precipitations which often serves as a point of failure and loss of RNA materials. In uPRO, we replaced it with direct buffer exchange between the consecutive affinity purifications and a column-based RNA purification to further shorten time and improve RNA yield.

PRO-seq requires two 5' RNA end pair chemistries: de-capping and phosphorylation to modify the 5' ends to become acceptor sites for RNA adaptor ligation. In uPRO, we proceed directly to reverse transcription using template switching to produce cDNA and add 5' adaptor sequence at the same time[28]. The cDNA product is processed through SPRI bead clean-up steps which remove short unused excess adaptors and primers. This serves as an additional enrichment step that reduces unwanted adaptor dimer products. As a result, the amount of amplified product in uPRO is more predictable than the conventional PRO-seq. Also, uPRO does not require test amplifications or polyacrylamide gel electrophoresis (PAGE) purifications (**Fig S1B**). Adaptor dimer product is relatively less and usually negligible for the Illumina sequencing (**Fig S1C**). Overall, uPRO may take as short as 6.5 hours to complete, compared to the 4-day conventional PRO-seq procedure (**Fig S1A**).

***Development of an artificial neural network program for bidirectional transcription scan, deepBTS.*** To detect bidirectional transcription signature of enhancers and promoters from uPRO data, we devised an artificial neural network deepBTS (**Fig 2A**). DeepBTS can predict TSSs from uPRO or PRO-seq data, make training sets from reference TSSs, and train neural networks with various designs.

To vectorize nascent transcription profiles as inputs for neural network analysis, we binned uPRO/PRO-seq reads along the genomic coordinates. We reasoned that multi-layer binning would capture the bidirectional signature efficiently through the combination of focused window in high resolution, wide range window in lower resolution, and/or intermediate range/resolution windows. We specifically tested two combinations of multi-layer binning (**Fig S2C**): 1) 50 bp high resolution bins in 500 bp region combined with 500 bp lower resolution bins in 5 kb regions; 2) Combination of 25 bp bin in 250 bp region, 50 bp bin in 1.5 kb region, and 500 bp bin in 10 kb region. The read counts on both strands are pseudo-logistic transformed to continuous variable vectors between 0 and 1. This vectorization is performed for all transcribed regions (more than 3 reads in 10 kb window) in 50 bp or 25 bp steps in uPRO/PRO-seq datasets.

For the output labels in the training sets, we used the previous PRO-cap defined eRNA and gene TSS references in LCLs. Random non-TSS coordinates used as non-TSS labels. We made use of the 16 PRO-seq data in 10 LCL cell lines (each replicated) for the training sets (14 datasets) and the test sets (GM19238, GM19239). The vectorized PRO-seq inputs and TSS output were used as the training set vectors. DeepBTS was coded in C++ using the FANN (Fast Artificial Neural Network) library. The network is trained with the resilient backpropagation (RPROP) algorithm for the 1’st pass to detect the network with the lowest error, then continued with the standard back propagation (SBP) algorithm to further reduce the error (**Fig S2A**).

The training can be performed using various neural network designs, such as number of hidden layers, number of neurons in each layer, and neuron connection properties. In deepBTS, the number of input neurons is determined by the binning options. For example, type 1 binning option leads to 80 input neurons ( 500 bp upstream and downstream in 50 bp bins (20 bins) + 5 kb up/downstream in 500 bp bins (20 bins) = 40 bins, in both plus and minus strands). Number of output neurons is 2, either a bidirectional TSS (BTS) or non-BTS. For the number of hidden neurons, we only tested the nearest integers of the geometric progression from the number of input to the number of output (e.g. 80, 12, 2 or 80, 24, 6, 2), depending on the number of hidden layers (**Fig S2C**). The variations of number of neurons around this range did not produce significant differences in the training performances.

After training the networks, we evaluated the performance using the 2 test datasets that were not used in the training (GM19238, GM19239). Using the PRO-cap based LCL TSS reference as the gold standard, we calculated true positive and false positive rates at different deepBTS score cut-off thresholds and generated AUROC plots (**Fig 2B, S2B**). We saw improvement of AUROC through a trial and error approach to find better binning and training parameters. Notably, switching algorithms produced better performance (**Fig S2B;** 608-1 to 608-2; 614-1 to 614-2). However, AUROC decreases after consecutive retraining, showing signs of overfitting (**Fig S2B;** 419-1 to 419-2; 614-2 to 614-3). We found that initial training with the RPROP algorithm to find the local minimum of error, then one standard back-propagation re-training yielded the best result in terms of AUROC.

***Cell type decomposition using comparative analysis between mRNA expression data and pChRO.***
Whereas the distinction between PBMC and PMNL is the biggest portion of peripheral leukocyte fractions, subpopulation compositions within PBMC can also influence the overall pChRO profile. Since PBMCs have much higher transcriptional activity than PMNL, PBMC subpopulations such as B cell and T cell subtypes can also influence the overall pChRO expression profile. To further investigate this possibility and subclassify PBMC populations, we performed a comparative analysis between existing RNA expression data in PBMC subtypes[26] and our pChRO data.

We first compared our uPRO data from PBMCs and PMNLs to a microarray data in all leukocytes: PBMC, B cell, T cells, and CD8+ T cells. Direct comparison between nascent RNA sequencing and mRNA microarray results can be affected by a lot of variables which does not make it feasible. Instead, we normalized the cell type specific uPRO data by all leukocyte pChRO profiles and microarray subtype data by all leukocyte results. We saw that there is significant global correlation between uPRO PBMC and microarray PBMC when normalized by all leukocyte data (**Fig S7**). In addition, microarray data in other PBMC subtypes such as B cells and CD4+ T cells are correlated with the uPRO PBMC data. On the other hand, uPRO PMNL data did not show any significant correlation with any other microarray-based cell subtype data. Normalizing the microarray data by cell types other than all leukocytes made the correlations disappear, indicating that the correlation between uPRO PBMC and microarray PBMC cell subtypes are specific to the cell subtype and proper normalization (**Fig S7**).

After confirming that the nascent RNA sequencing data is in agreement with the existing mRNA expression data, we compared the signature gene lists from both data sets. The PBMC signature genes from the uPRO data are driven mostly by strong depletion in the PMNL population (**Fig S7**). According to the mRNA data, enrichment of the signature genes in PBMC subpopulations was variable. However, we identified groups of the uPRO PBMC cluster genes that appear more specific to B cell or T cell subpopulations. On the other hand, PMNL signature genes that are depleted in PBMC cells also appear depleted in the mRNA microarray data (**Fig S7**).

Conversely, the mRNA microarray signature genes show expression level differences in the uPRO data (**Fig S7**). The mRNA data did not include PMNL cell isolates as an exclusive comparison but rather used the subtractive enrichment or depletion in PBMC over all leukocyte as a PBMC marker (labeled LYMPHS) or PMNL marker (labeled GRANS). Therefore the PBMC(LYMPHS, green) signature gene expression was not as prominent in the uPRO PBMC data (pPBMC). However other signature genes in B, T, and CD8 cells are more strongly enriched (**Fig S7**). Conversely, the PMNL (GRANS, blue) signature gene expression is markedly reduced in PBMC showing the reproducibility of mRNA microarray signature genes in uPRO data.

We compared the individual pChRO data to the mRNA microarray signature genes (**Fig S7**). Consistent with the uPRO signature genes (**Fig 7G**), we saw an overall increase in PMNL (GRANS) signature genes in Individual 2 which indicates an increase in the PMNL subpopulation (**Fig S7**). Overall, these results demonstrate that nascent RNA sequencing and mRNA microarray are in good agreement and can be used together to further deconvolute specific cell subtypes from the pChRO data.

### Supplementary Figures

#### Figure S1

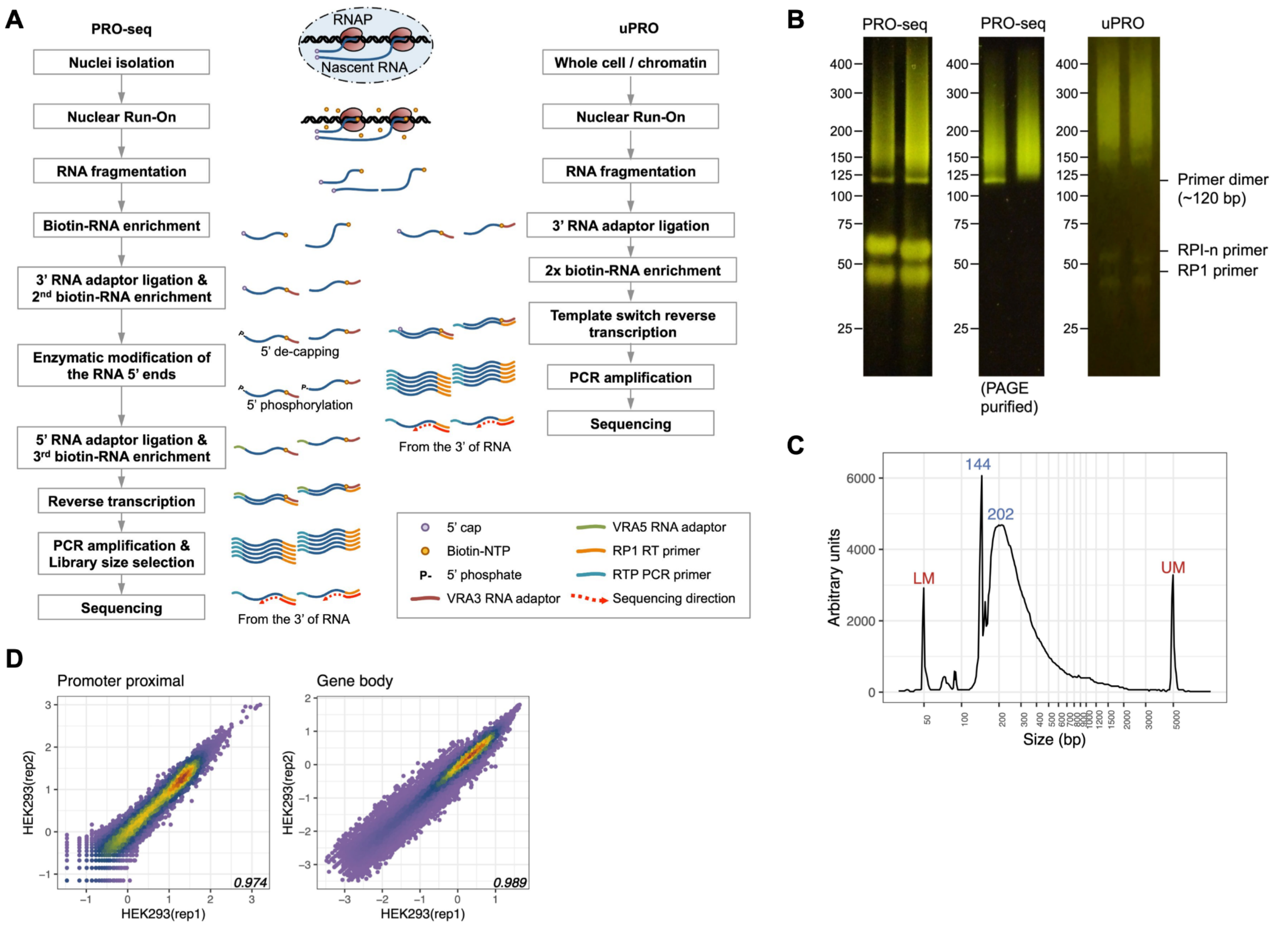


**Fig S1.** Schematics of the uPRO procedure. **A.** Comparison between conventional PRO-seq and uPRO procedures (PRO-seq procedure adapted from Mahat et al[12]). **B.** Polyacrylamide gel electrophoresis of PRO-seq and uPRO libraries. **C.** Capillary electrophoresis trace (BioAnalyzer) of a representative uPRO library. LM: lower marker (50 bp), UM: upper marker (5,000 bp). **D.** Correlation scatterplot of PRO-seq replicates in HEK293 cells.

**Figure S2**

A

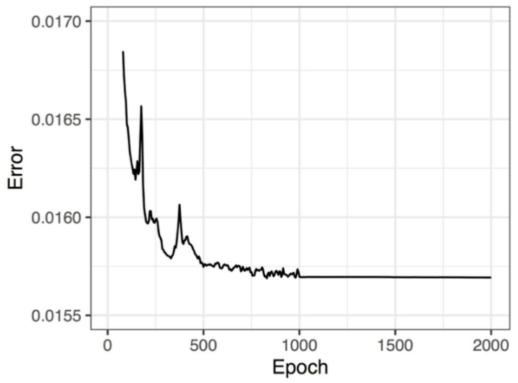


C

| Network | # hidden | Binning type | Training set | Initial training | Re-training | Note |
|---|---|---|---|---|---|---|
| 327 | 1 | 2 | GM18505 | SBP | - | 2000 epochs, 27 hr 24 min |
| 330 | 1 | 1 | GM18505 | SBP | - | 1000 epochs, 1 hr 56 min |
| 419 | 1 | 1 | GM18505 | SBP | SBP | |
| 429 | 1 | 1 | All | SBP | SBP | |
| 607 | 1 | 1 | All | RPROP | SBP | Used in the paper |
| 608 | 1 | 1 | All | RPROP | SBP | Repeat of 607 |
| 614 | 1 | 2 | All | RPROP | SBP + SBP | |
| 616 | 2 | 2 | All | RPROP | SBP | |
| 701 | 1 | 1 | GM19238r, GM19239r | RPROP | SBP | |

B

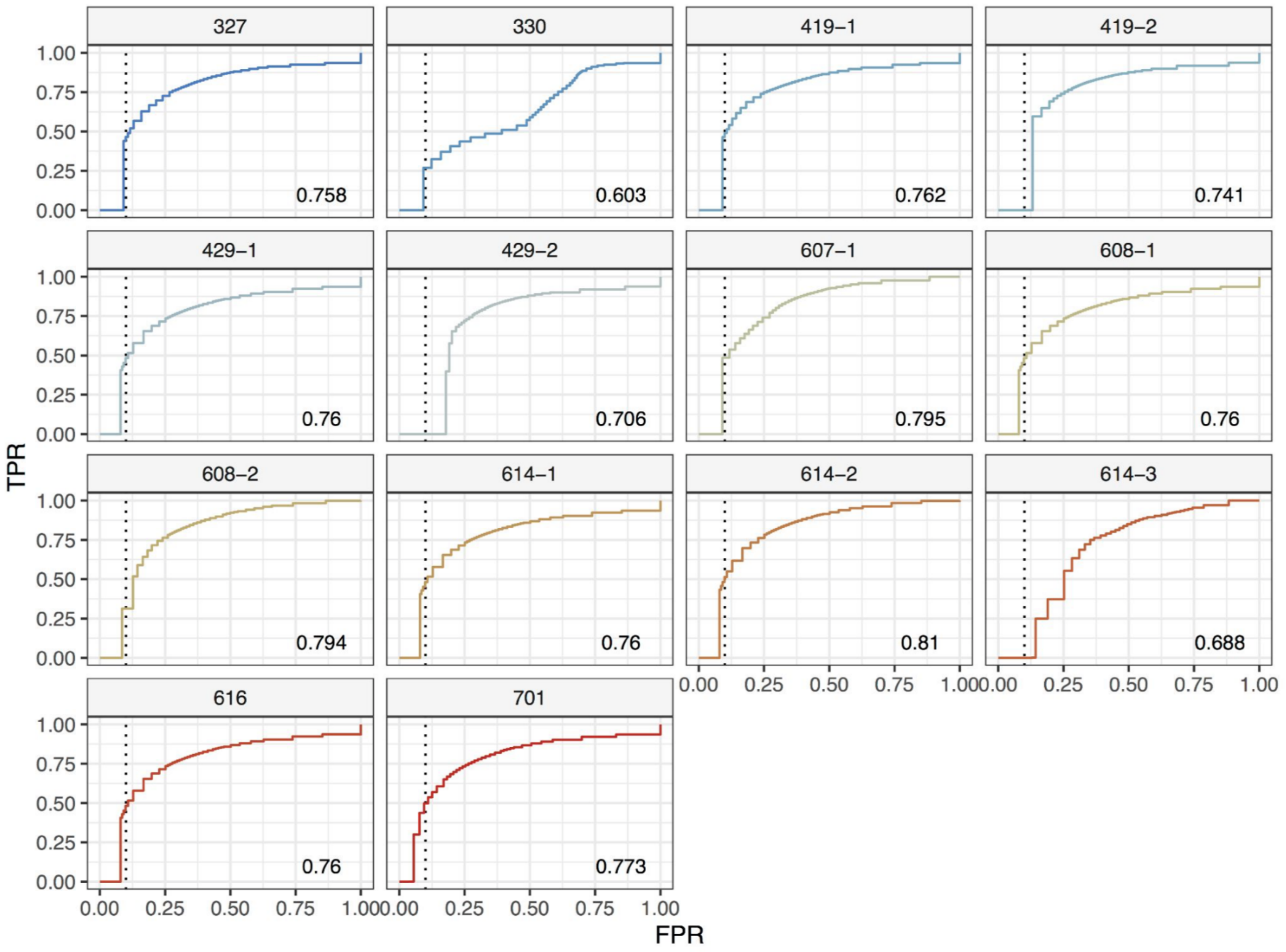


**Fig S2.** Deep learning based bidirectional transcription scan (deepBTS). **A.** A typical training curve or error per epoch (training round) for the 607 network. **B.** Receiver-operating response curve (ROC) of the trained networks. AUROC values at the lower right parts of the panels. **C.** Table of network training parameters.

**Figure S3**

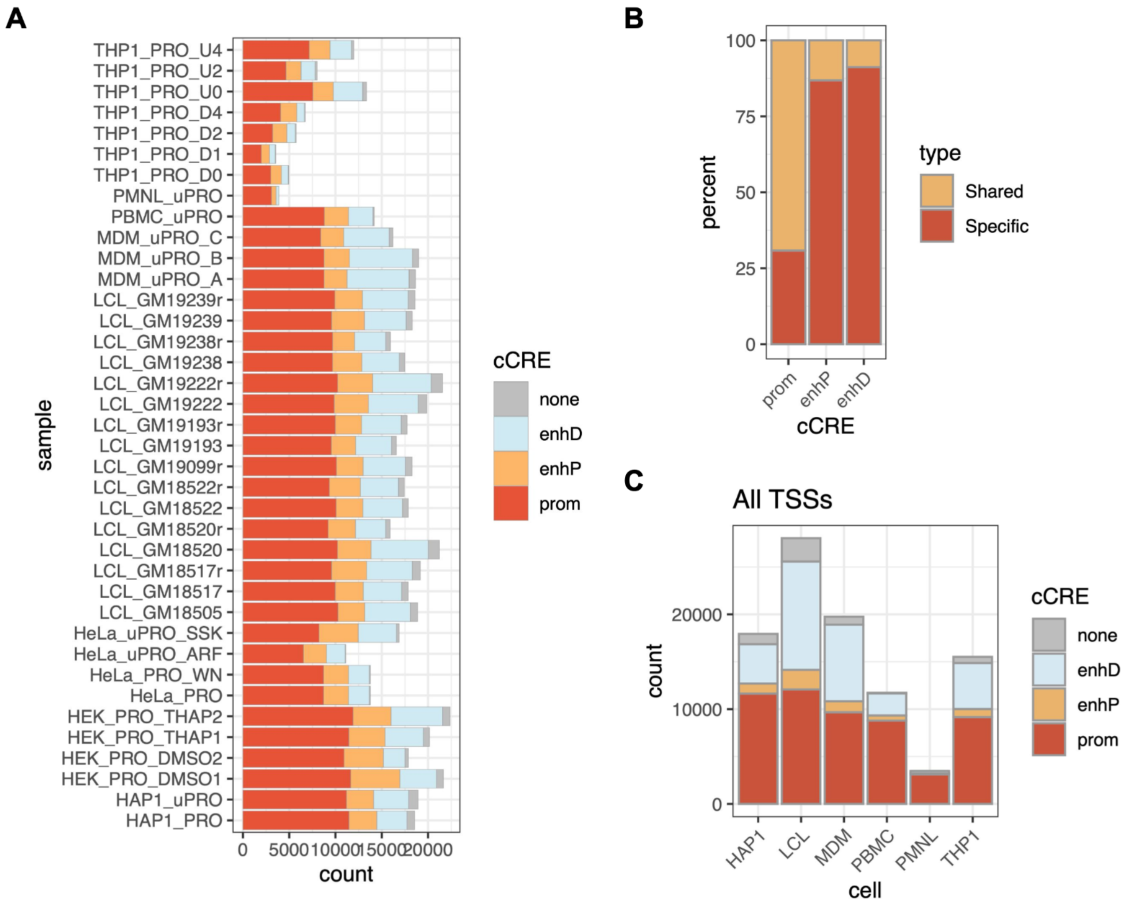


**Fig S3.** Identification of cell type specific gene and enhancer TSSs. **A.** Number of deepBTS TSSs categorized by ENCODE cCRE in each dataset. **B.** Proportions of deepBTS TSSs that are conserved (shared) and cell type-specific within blood cell types, categorized by ENCODE cCRE. **C.** Number of all TSSs combined for each cell type.

**Figure S4**

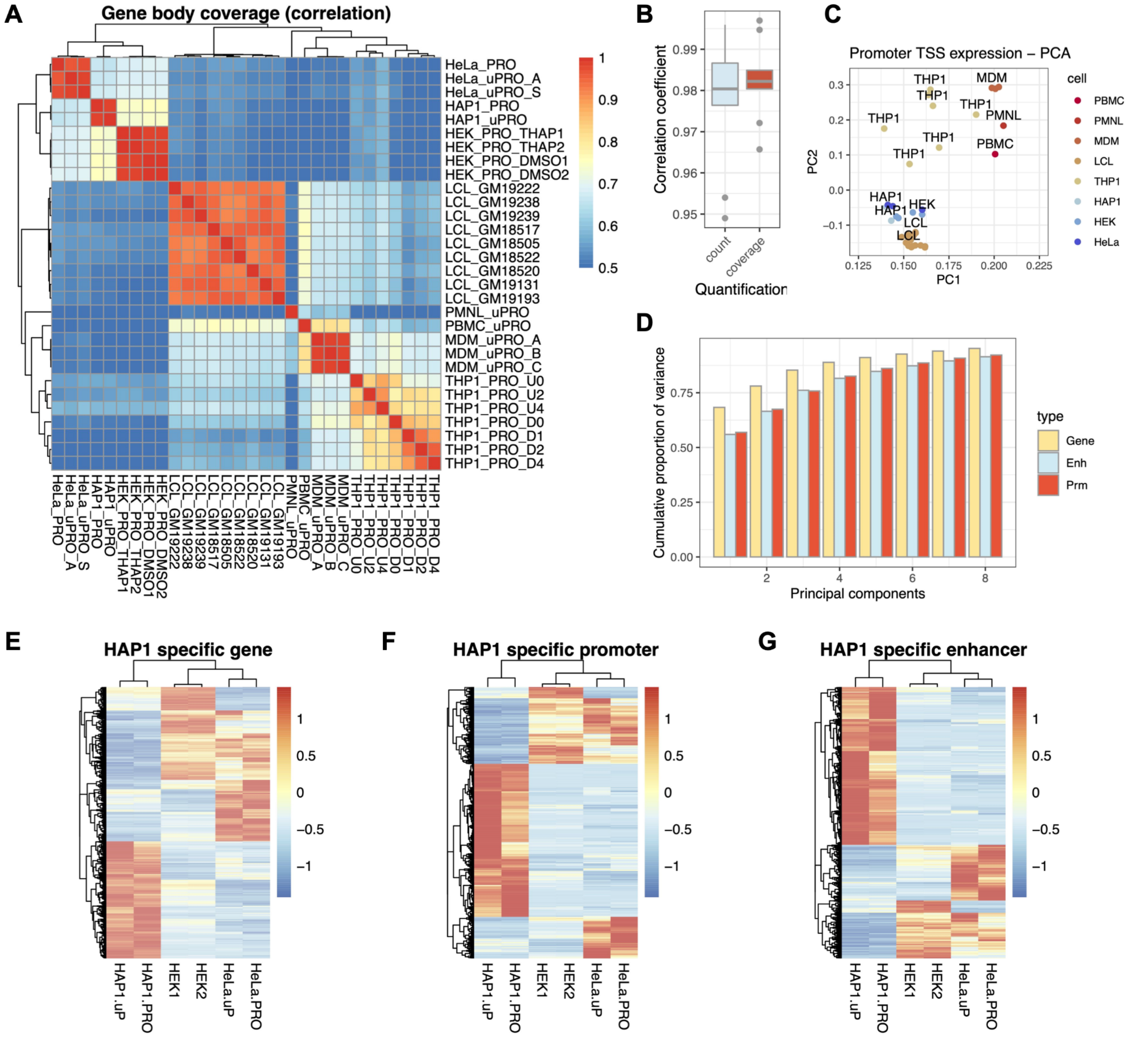


**Fig S4.** Identification of cell type specific genes and non-coding RNA expression. **A.** Heatmap of correlation coefficients between gene body read coverage of the datasets. **B.** Correlation coefficients between replicated datasets depending on the quantification method. **C.** Principal component biplot of promoter TSS expression. **D.** Principal component cumulative variance plots for gene, enhancer, and promoter expression. **E.** Clustered heatmap of HAP1 specific gene expression. Color label represents $\log_2$ fold difference from the mean of all cell types. Column labels suffixes are .u: uPRO, .pro: PRO-seq, 1: PRO-seq replicate 1, 2: PRO-seq replicate 2. **F.** Heatmap of HAP1 specific dBTS promoters. **G.** Heatmap of HAP1 specific dBTS enhancers.

**Figure S5**

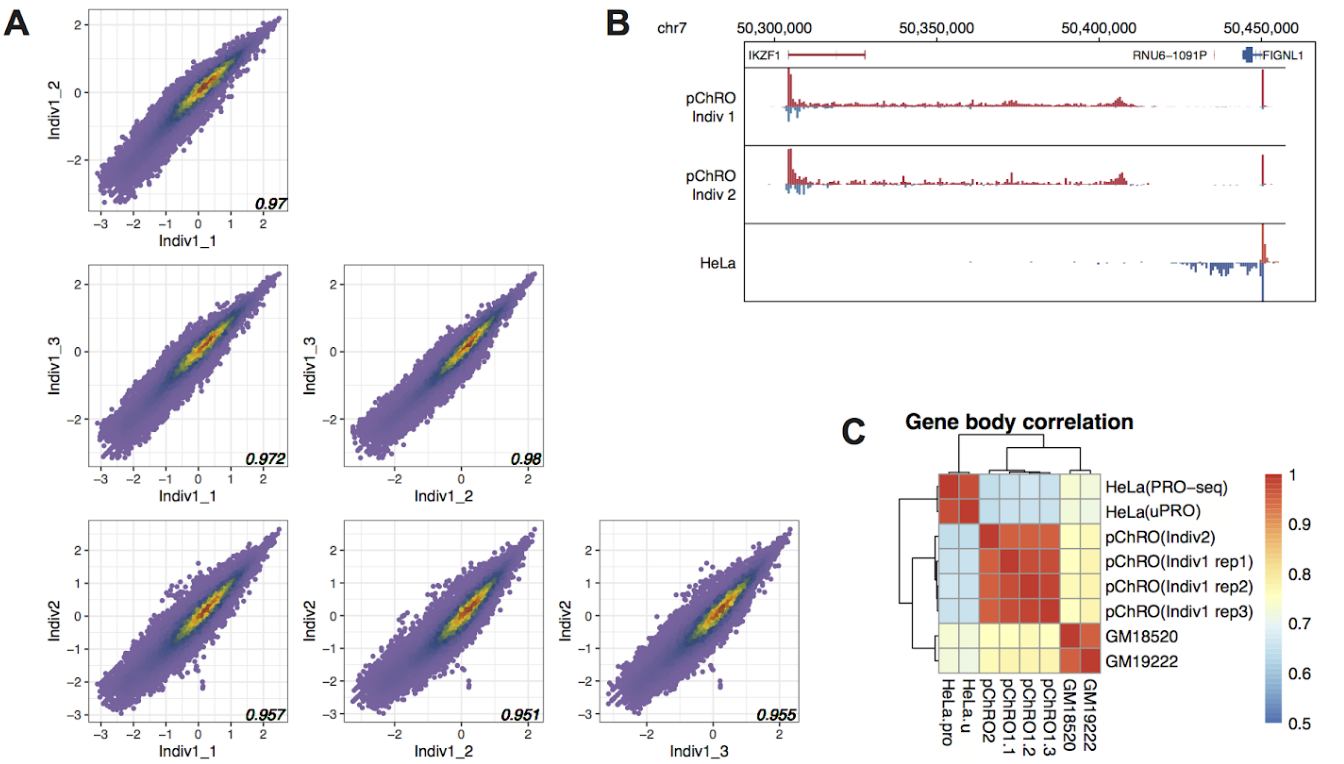


**Fig S5.** Direct ChRO-seq in peripheral blood samples (pChRO). **A.** Scatterplots of gene body correlations in replicates and different individuals. **B.** Example browser view of peripheral blood specific transcription. **C.** Heatmap of correlation clustering in pChRO and other PRO-seq data

**Figure S6**

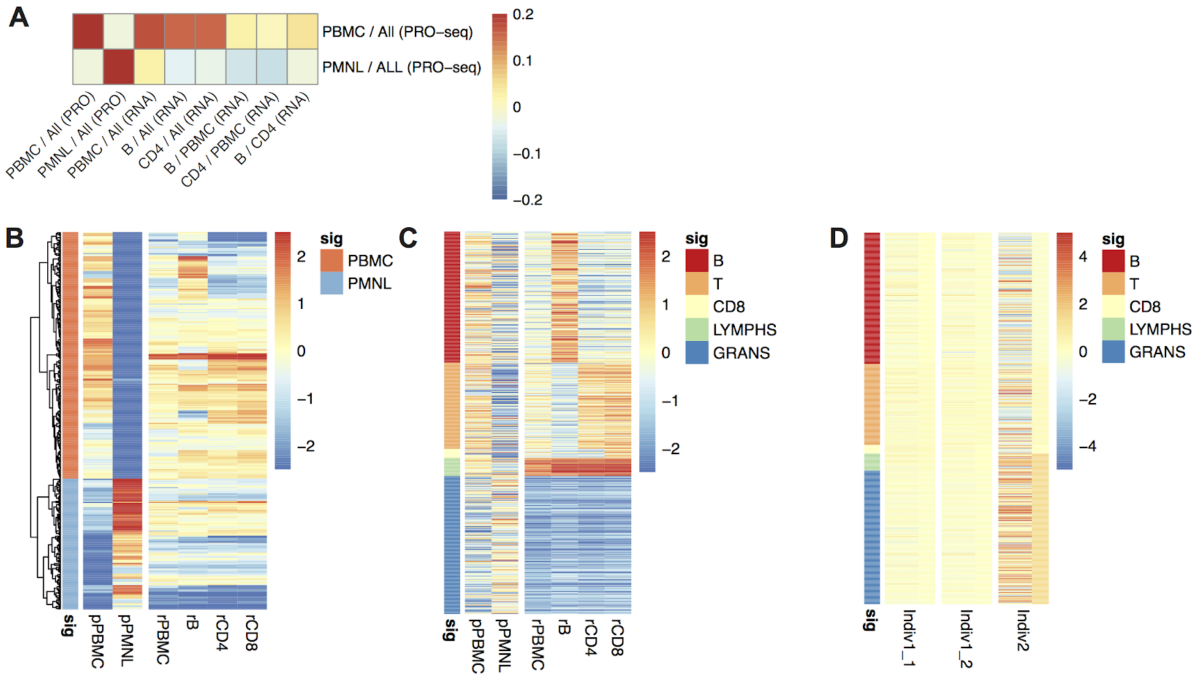


**Fig S6.** Decomposition of RNA-seq based sPBMC subtypes and identification of differentially expressed genes. **A.** Correlation heatmap between PRO-seq and mRNA microarray data from different peripheral leukocyte subpopulations. Color scale bar: Pearson correlation coefficient of the $\log_2$ read count/microarray levels normalized to the reference cell type indicated on the denominator. **B.** Heatmap of PRO-seq PBMC and PMNL signature gene expressions in PRO-seq or mRNA microarray data in different subpopulations. Column prefixes: p-PRO-seq, r-mRNA microarray. Color scale bar: $\log_2$ fold difference. **C.** Heatmap of microarray leukocyte subtypes (B, T, CD8+ T, LYMPHS = PBMC, GRANS = PMNL). signature gene expressions in PRO-seq or mRNA microarray data. Column prefixes: p-PRO-seq, r-mRNA microarray. Color scale bar: $\log_2$ fold difference. **D.** Heatmap of microarray signature gene expression (B, T, CD8+ T, LYMPHS = PBMC, GRANS = PMNL) in pChRO individual samples. Thin ribbons on the right represent the average fold difference of the signature gene group in the corresponding individual. Color scale bar: $\log_2$ fold difference.

### Supplementary Tables

**Table S1.** List of deepBTS TSS regions, and classification

**Table S2.** Normalized read counts in deepBTS regions

**Table S3.** Normalized read coverage at gene bodies in cell lines

**Table S4.** Hierarchical co-expression network in cell lines

**Table S5.** Table of PBMC and PMNL signature genes
